# Chronic morphine administration differentially modulates viral reservoirs in SIVmac251 infected rhesus macaque model

**DOI:** 10.1101/2020.09.09.290700

**Authors:** Arpan Acharya, Omalla A Olwenyi, Michellie Thurman, Kabita Pandey, Brenda M Morsey, Benjamin Lamberty, Natasha Ferguson, Shannon Callen, Qiu Fang, Shilpa J. Buch, Howard S. Fox, Siddappa N. Byrareddy

**Author notes:** Corresponding author: Siddappa N. Byrareddy, (SNB). These authors contributed equally to this work (Shilpa J. Buch, Howard S. Fox, Siddappa N. Byrareddy).

## Abstract

HIV persists in cellular reservoirs despite effective anti-retroviral therapy with rebound of viremia upon therapy interruption. Opioids modulate the immune system and suppress antiviral gene responses, which significantly impact people living with HIV (PLWH). However, the effects of opioids on viral reservoir remains elusive. Herein, we describe a morphine dependent SIVmac251 infected Rhesus macaque (RM) model to study the impact of opioids on HIV reservoirs. RMs were ramped up with morphine (n=10) or saline (n=9) for two weeks to a final dosage of 5mg/kg administered twice daily, which was maintained for seven weeks, and then infected with SIVmac251. Combined anti-retroviral therapy (cART) was initiated in approximately half the animals in each group five weeks post-infection and morphine/saline administration continued for 10 months. Among drug naïve macaques, there were no differences in plasma/CSF viral load nor in cell-associated DNA/RNA loads. However, within the cART-suppressed macaques, there was a reduction in cell-associated DNA load, intact proviral DNA copy numbers, and inducible SIV reservoir in both peripheral blood and lymph nodes (LNs) of morphine-administered RMs compared to saline controls. Further, PBMCs of morphine administered RMs, a reduction in Th1 polarized CD4+ T cells and in LNs there was a reduction in the total Tfh and Th1 like Tfh cells were observed, indicating probably have impact on reduction of viral reservoirs. In distinction to PBMC and LNs, within the CNS size of latent SIV reservoirs was higher in the CD11b+ microglia/macrophages of morphine-dependent RMs. These data suggest that morphine plays a role in modulating SIV reservoirs, reduces the CD4+ T-cell reservoir in both peripheral blood and LNs, and increases microglia/macrophage reservoirs in CNS. These findings will aid in understanding of molecular mechanism(s) of opioid-mediated differential modulation of viral reservoirs and evaluation of therapeutic strategies to reduce/eliminate HIV reservoirs in opioid-dependent PLWH.

**Author summary:** Opioids are commonly used as well as abused by HIV infected individuals, are known to suppress immune responses. However, their effects on modulating viral reservoir dynamics is not known. Here we developed a morphine dependent SIVmac251 infected rhesus macaque (RM) model to study the impact of opioids on HIV reservoirs and immune cells. We found that there was no difference in viral loads or cell-associated DNA/RNA loads among morphine dependent vs. saline treated control macaques. On the other hand, when macaques were treated with cART, there was a reduction in SIV reservoirs both in periphery and lymphoid tissues in morphine administered RMs, and the size of latent SIV reservoir was higher in the CNS CD11b+ microglia/macrophages as compared to control macaques. Therefore, these new data form macaque models suggest that PLWH who suffering from opioid use disorders have higher reservoirs in CNS as compared to lymphoid system. Thus, through understanding these reservoirs among PLWH who uses opioids are critical for better designing HIV cure strategies.

## Introduction

Opioid use disorder (OUD) is a chronic illness characterized by persistence use of opioids to the detriment of the user [1]. Globally an estimated 26.8 million people were living with OUD in 2016 [1], of which 2.1 million were from USA [2]. People living with OUD are at higher risk of medical comorbidities including human immunodeficiency virus (HIV) and hepatitis C (HCV) infection [3]. Globally it is estimated 80% of injection drug users take opioids and 17.8% of them are HIV positive [4, 5]. Epidemiological studies have shown that chronic opioid administration is immunosuppressive, downregulates the antiviral genes and increases the susceptibility to infections like tuberculosis and HIV [6-12]. The innate and adaptive immune responses act in synchrony against the invading pathogens, and opioids acting through the opioid receptors alter both these responses [13, 14].

OUD is important in the context of HIV infection, as several studies have reported that PLWH who use opioids are at a higher risk of developing neurological disorders compared to PLWH without OUD [15-20]. Opioids have been shown to upregulate the expression of CCR5 while downregulating the expression of its cognate β-chemokine production by acting through µ-opioid receptor in macrophages and microglia, thereby boosting the entry of R5 tropic viruses in these target cells [21-24]. Opioids and HIV synergistically act to activate glia and upregulate the expression of cytokines and chemokines, leading, in turn, to neuronal damage and development of hyperalgesia [25, 26].

One barrier to the cure of HIV is that the HIV genome integrates into the host chromosome [27]. A subset of infected cells with integrated proviral DNA can enter a transcriptionally silent state that can persist in the context of combination antiretroviral therapy (cART) and escape host cell immune surveillance owing to its inability to produce antigens essential for triggering immune responses. This, in turn leads to creation of a pool of long-lived viral reservoirs [28]. HIV reservoirs are seeded early after infection and persist through the entire lifetime of the host. Although among all the cellular reservoirs, resting memory CD4+ T cells are the best characterized, persistence of viral reservoirs in perivascular macrophages and microglia in CNS has also been reported [29, 30]. Active and latent reservoirs of HIV also persist in several other tissues owing to the presence of subtherapeutic concentrations of antiretrovirals in these tissue locations. The major barrier to penetration of ART in deep tissues are their physiochemical properties and the presence of drug efflux transporters [31]. Among the well-characterized tissue reservoirs such as LNs, spleen, gut associated lymphoid tissues (GALT) and the thymus [32]. The other less characterized anatomical tissue reservoirs of HIV include tissues such as the kidney, liver, lung, bone marrow, genital tract and CNS [33-35].

Morphine and other frequently used/abused opioids have been shown to inhibit phagocytic property of macrophages, migration of neutrophils and have a cytopathic effect on NK cells, which in combination results in dampening of the innate immune system [36]. In adaptive immune responses, chronic morphine exposure leads to defective antigen presentation and a phenotype switch from Th1 to Th2 in CD4+ T cells resulting in inhibition of pro-inflammatory responses that are critical for eradicating invading intracellular pathogens [37]. Interestingly, there are also conflicting reports refuting the immune suppressive effects of morphine, with some reports from rodent models and studies using ex-vivo human samples indicating activation of the pattern recognition receptor TLR4 and subsequent cellular activation [38-42].

There are multiple lines of evidence implicating the role of morphine in mediating immunosuppressive effects in the periphery, its role in infectivity, transmission and pathogenesis of HIV and its potentiation of HIV neuropathogenesis [16, 21, 24, 43, 44]. However, role of morphine in modulating the persistence of HIV reservoirs in various anatomical sanctuaries, remains elusive. In the present study, we sought to examine the impact of morphine dependency on the size of SIV reservoirs using the SIVmac251-infected rhesus macaque (RM) model. SIV reservoirs were found to persist in all the major cellular and tissue locations in ART-suppressed, morphine-dependent, and saline controls. Our findings suggest that morphine differentially modulate of the size of viral reservoirs in periphery and lymphoid tissues versus the brain. To best of our knowledge, this is the first study to report differential regulation of viral reservoir dynamics in the context of opioid dependence and warrants further investigation to dissect the molecular mechanism(s).

## Results

### Dynamics of plasma and CSF viral loads in morphine dependent versus control SIVmac251 infected rhesus macaques with and without antiretroviral therapy

In this study, we included a total 19 macaques. Ten (n=10) RM were ramped up over two weeks to a final dose of twice daily 5 mg/kg dose of morphine, which was maintained for seven weeks, while nine (n=9) RMs received saline, that served as control. At week 9, all the macaques were inoculated with single 200 TCID_50_ dose of SIVmac251 by intravenous injection. Five weeks post-inoculation cART was initiated in six animals out of ten RM in the morphine group and in five out of nine RMs in the control group. Four monkeys from each group were left untreated till the end of the study (**S1 Fig**). Within the control group (n=9) the median peak plasma and CSF viral load was 1.25×10^7^ SIV copies/ml and 1.02×10^5^ SIV copies/ml respectively. On the other hand, in the morphine administered group (n=10), median peak plasma and CSF viral load was 8.68×10^7^ SIV copies/ml and 4.62×10^4^ SIV copies/ml (p=0.447 for plasma and p=0.400 for CSF) (**Fig 1A, B, E, F**). All eleven animals from the antiretroviral treated group became aviremic with completely suppressed plasma and CSF viral load within 13 weeks post-inoculation which is equivalent to 8 weeks of antiretroviral therapy and they remained virally suppressed till the end of this study (**Fig 1E, F**). There were no statistical differences either peak plasma/CSF viral loads or longitudinal geometric mean of plasma/CSF viral loads between morphine vs. saline administered RMs in all the groups (**Fig 1 C, D, G, H**).

**Fig 1:**
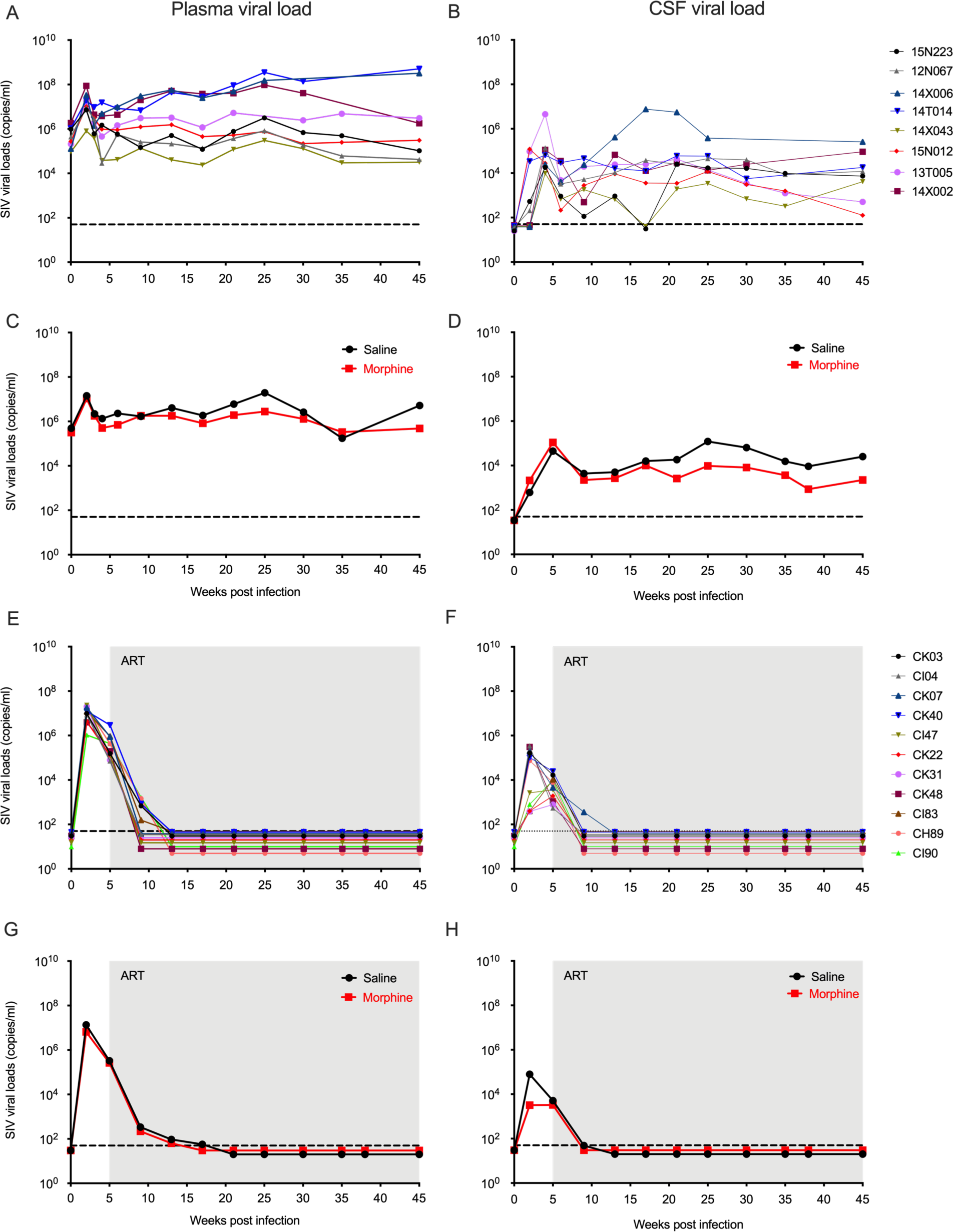
Plasma and CSF viral load of morphine administered SIVmac251 infected *rhesus macaques*. Ten macaques were ramp-up with daily twice dose of morphine (5 mg/kg) for eight weeks and nine macaques were served as control (received saline), then infected with SIVmac251. Five weeks post infection, in six animals from morphine group and five animals from control group daily single dose of combination antiretroviral therapy (cART) was initiated. Daily dose of cART and morphine/saline administration continued until the end of the study. Plasma and CSF SIV viral load was measured longitudinally in all *rhesus macaques* using quantitative PCR (QPCR) assay. **(A)** Longitudinal plasma viral loads of individual animals that did not receive cART (copies/ml); **(B)** Longitudinal CSF viral loads of individual animals that did not received cART (copies/ml); **(C)** Geometric mean of plasma viral loads of animals that did not received cART (black line represents control group who received saline and red line represents morphine treated group); **(D)** Geometric mean of CSF viral loads animals that did not received cART (black line represents control group who received saline and red line represents morphine treated group). **(E)** Longitudinal plasma viral loads of individual animals received cART (copies/ml); **(F)** Longitudinal CSF viral loads of individual animals received cART (copies/ml); **(G)** Geometric mean of plasma viral loads of animals received cART (black line represents control group who received saline and red line represents morphine treated group); **(H)** Geometric mean of CSF viral loads animals received cART (black line represents control group who received saline and red line represents morphine treated group). The shaded region indicates ART phase of the study. The dashed lines represent limit of detection of the assay (50 copies/ml).

### Cell-associated SIV DNA/RNA in the blood, lymph nodes and rectal mucosal tissues

To understand how chronic morphine administration could modulates the size of SIV reservoirs in various tissue compartments in the presence and or absence of cART, we sought to quantify the cell-associated SIV DNA and RNA in various anatomical tissue sanctuaries collected during necropsy of all the RMs included in this study. Since CD4+ T cells are known to be the major contributor of viral reservoirs in both the blood and LNs, CD4+ T cells were purified from peripheral blood and LNs followed by isolation of DNA/RNA as described in Material and Methods. In ART naïve RMs, in the PBMC the median cell-associated SIV DNA load was 15,898 copies/10^6^ CD4+ T cells in morphine administered group versus 8,547 copies/10^6^ CD4+ T cells in saline controls (p=0.600). In ART treated group, in PBMC the median cell-associated SIV DNA load was 3,332 copies/10^6^ CD4+ T cells in morphine administered group versus 4,583 copies/10^6^ CD4+ T cells in saline controls (p=0.055) (**Fig 2A**). Similarly, in ART naïve RMs, in PBMC median cell-associated SIV RNA load was 30,533 copies/10^6^ CD4+ T cells in morphine administered group versus 18,890 copies/10^6^ CD4+ T cells in saline controls (p=0.990). In the ART treated group, in PBMC the median cell-associated SIV RNA load was 94 copies/10^6^ CD4+ T cells in morphine administered group versus 446 copies/10^6^ CD4+ T cells in controls (p=0.120) (**Fig 2B**).

**Fig 2:**
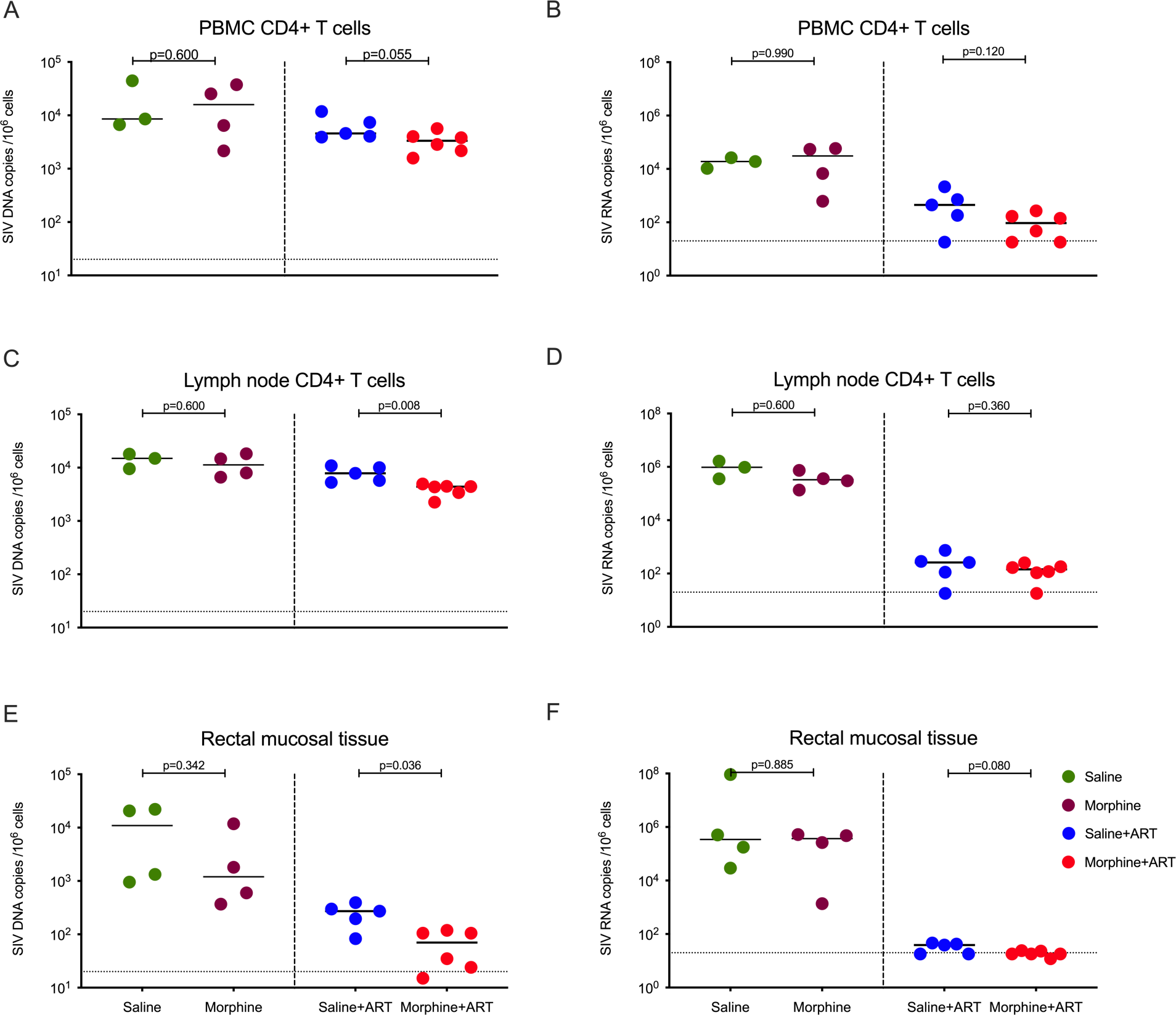
Cell associated SIV DNA and RNA level in different tissue compartments between morphine administered versus control groups of SIVmac251 infected *rhesus maca*ques. **(A)** SIV DNA copies per million CD4+ T cells isolated from peripheral blood mononuclear cells; **(B)** SIV RNA copies per million CD4+ T cells isolated from peripheral blood mononuclear cells; (**C)** SIV DNA copies per million CD4+ T cells isolated from lymph node (LN) cells; **(D)** SIV RNA copies per million CD4+ T cells isolated from LN cells; **(E)** SIV DNA copies per million cells from rectal mucosal tissue; **(F)** SIV RNA copies per million cells from rectal mucosal tissue; The green dots represent samples from control animals administered saline that did not received ART, maroon dots represent samples from morphine administered animals that did not received ART, blue dots represent samples from control animals administered saline and treated with ART and the red dots represent samples from morphine administered animals treated with ART. The horizontal dashed lines represent limit of detection of the assay (20 copies/ml). All values below limit of detection are bringing to the limit of detection for better representation in the figure.

Next, we measured the cell-associated DNA and RNA in CD4+ T cells purified from LNs. In LNs of ART naïve RMs, median cell-associated SIV DNA load was 11,295 copies/10^6^ CD4+ T cells in morphine administered group versus 14,962 copies/10^6^ CD4+ T cells in saline controls (p=0.600). On the other hand, in LNs of the ART group a significant difference was found, with a median cell-associated SIV DNA load of 4,392 copies/10^6^ CD4+ T cells in morphine administered group versus 7,847 copies/10^6^ CD4+ T cells in saline controls (p=0.008) (**Fig 2C**). The RNA loads did not significantly differ. In LNs of ART naïve RM, median cell-associated SIV RNA was 330,802 copies/10^6^ CD4+ T cells in morphine administered group versus 968,610 copies/10^6^ CD4+ T cells in saline controls (p=0.220). In the LNs of ART treated group, median cell-associated SIV RNA load was 143 copies/10^6^ CD4+ T cells in morphine administered group versus 260 copies/10^6^ CD4+ T cells in saline controls (p=0.360) (**Fig 2D**).

Further, we assessed cell-associated DNA and RNA in mucosal tissue collected from the rectum during necropsy. In the ART naïve RMs median cell-associated SIV DNA load was 1,201 copies/10^6^ cells in morphine administered group versus 10,961 copies/10^6^ cells in saline controls (p=0.342). In the ART treated group, a significant difference was found, with a median cell-associated SIV DNA load of 70 copies/10^6^ cells in morphine administered group versus 272 copies/10^6^ cells in saline controls (p=0.036) (**Fig 2E**). In ART naïve RMs median cell-associated SIV RNA load was 368,990 copies/10^6^ cells in morphine administered group versus 341,883 copies/10^6^ cells in saline controls (p=0.885). In the ART treated group, the median cell-associated SIV RNA load was 11 copies/10^6^ cells in morphine administered group versus 39 copies/10^6^ cells in saline controls (p=0.080) (**Fig 2F**).

We also measured the cell-associated DNA/RNA in both the spleen and lungs of macaques. There was no statistically significant difference in cell-associated SIV DNA level in the morphine administered group versus controls in presence of ART treatment in both spleen (p=0.320) and lung (p=0.990), as well as in in the spleen (p=0.890) or lungs (p=0.190) of ART naïve RMs (**S2A** and **S2C Fig**). Similarly, we did not observe any significant difference in cell-associated SIV RNA levels in the spleen (p=0.520) or lungs (p=0.920) of either morphine administered group versus saline controls in ART treated RMs or in the spleen (p=0.890) and lungs (p=0.310) of ART naïve morphine administered versus saline controls (**S2B** and **S2D Fig**).

In summary, we found a moderate reduction of tissue associated viral loads in both DNA and RNA levels in morphine administered cART treated animals as compared with saline controls.

### Quantitation of SIV reservoirs in CD4+ T cells of blood and lymph nodes

We used the Tat/rev Induced Limiting Dilution Assay (TILDA) to measure the size of inducible SIV reservoirs in the blood and LNs of morphine/saline administered, SIV-infected cART-treated RMs (**S1 Fig**). In peripheral blood, the median frequency of CD4+ T cells producing inducible multiply splices (ms)-Tat/rev transcript per million of CD4+ T cells was reduced, measuring 4.54 in morphine administered group versus 9.11 in the saline control group (p=0.067) (**Fig 3A)**. While that did not reach significance, in the LNs a similar reduction was found in the median frequency of CD4+ T cells producing inducible ms-Tat/rev transcript per million of CD4+ T cells as 6.47 in morphine administered group versus 10.38 in the saline control group which was statistically significant (p=0.036) (**Fig 3B)**.

**Fig 3:**
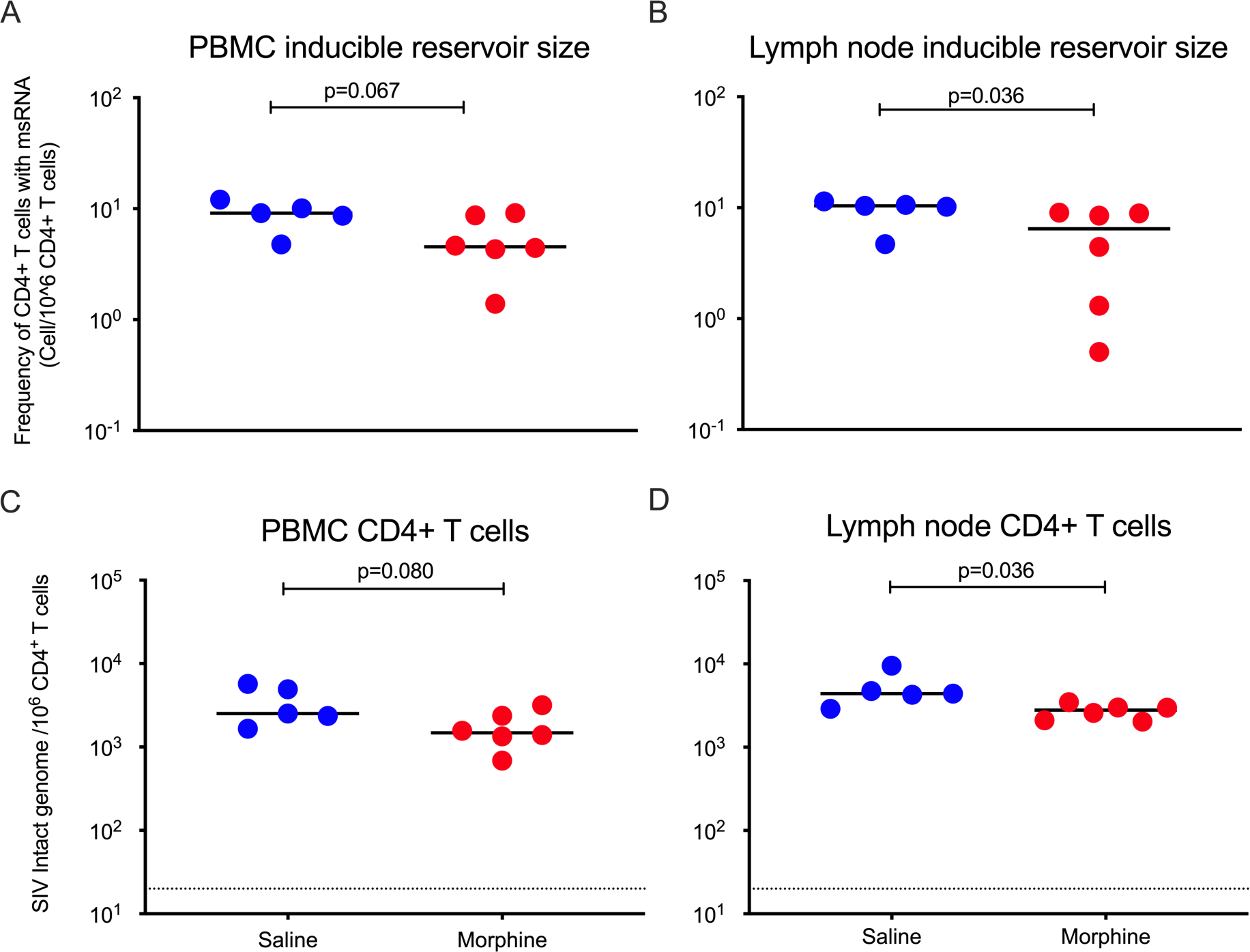
Size of SIV reservoirs in CD4+ T cells from peripheral blood and lymph nodes (LNs) in morphine administered versus control group of SIVmac251 infected ART suppressed *rhesus maca*ques. **(A, B)** Size of inducible SIV latent reservoirs in CD4+ T cells from peripheral blood and LNs of morphine administered and control rhesus macaques received saline were measured by Tat/rev Induced Limiting Dilution Assay (TILDA) which quantify frequency of CD4+ T cells producing inducible ms-Tat/rev transcript per million of CD4+ T cells. **(C, D)** Frequency of intact SIV genome in CD4+ T cells isolated from blood and LNs of morphine/saline administered, SIVmac251 infected cART treated rhesus macaques were quantified using IPDA.

We next used the intact proviral DNA assay (IPDA) to measure the size replication competent SIV reservoirs in both the blood and LNs. Recently Bruner et al., developed a multiplex droplet digital PCR (ddPCR) based HIV-1 proviral DNA quantification assay, that is able to distinguish between intact versus defective proviruses and is referred to as IPDA. The authors have described a positive correlation and identical decay rate of viral reservoirs when measured by quantitative viral outgrowth assay (QVOA) and IPDA [45]. Bender et al., have reported adaptation of IPDA for quantification of the intact SIV genome in ART suppressed rhesus macaques, implicating thereby its utility to serve as a surrogate marker to measure size of latent reservoirs [46]. Herein, using IPDA, we measured the frequency of intact SIV genome in both blood and LNs of morphine/saline administered as well as SIV-infected cART treated RM (**S1 Fig**). In CD4+ T cells purified from peripheral blood, the median of intact SIV genome per million cells was lowered by morphine, 1,480 for the morphine administered group versus 2,521 in the saline control group (p=0.080) (**Fig 3C**). While that did not reach significance, in the CD4+ T cells purified from LNs, morphine again lowered the median of intact SIV genome per million cells, measured at 2,789 for the morphine administered group versus 4,399 in saline control group, a statistically significant difference (p=0.036) (**Fig 3D**).

### Changes in CD4+ T cell polarization in PBMCs and Lymph nodes

Chronic morphine exposure depletes lymphoid cells and leads to a phenotypic switch of Th1 to Th2 in CD4+ T cells [37]. On the other hand, a recent report suggests that in chronic HIV patients, majority of intact replication competent proviruses persist in Th1 polarized CD4+ T cells [47]. To understand the impact of chronic morphine administration on T lymphocytes which have majority of HIV reservoirs in peripheral blood and LNs, at necropsy, flow cytometry was performed to determine the CD4+ T cell polarity in the PBMCs and LNs of cART treated RMs. In PBMC, to differentiate between Th1, Th2 and Th17 cells intracellular cytokine staining was performed as described earlier [48]. Due to poor cytokine production by LN germinal center Tfh cells, it is difficult to differentiate Tfh subpopulations by cytokine production assays [49]. Therefore, we performed surface staining of chemokine receptors of Tfh cells (CD95+PD1+CXCR5+) from LNs to identify Th1 like Tfh (CXCR3+), Th2 like Tfh (CCR4+) and Th17 like Tfh (CCR6+) subpopulations [50]. The gating strategies used for PBMCs and LNs is described in **S3, S4** and **S5 Fig**. In PBMCs, while not significantly different, morphine administered macaques had lower levels of Th1 polarized CD+ T cells compared to saline controls as denoted by lower levels of IFNγ and TNFα cytokine secreting cells with a median of 0.87% versus 1.55% (p=0.536) for IFNγ (**Fig 4A**) and with a median of 5.61% versus 14.70% for TNFα (p=0.017) (**Fig 4B**) respectively. The proportion of IL-4 secreting Th2 polarized CD4+ T cells were higher in the morphine administered macaques with a median of 2.55% versus 1.76% in saline controls (p=0.305) (**Fig 4C**). The levels IL-17 secreting CD4+ Th17 lymphocytes in the morphine treated group did not appreciable differ, with a median of 0.47% versus 0.48% in saline controls (p=0.930) (**Fig 4D**). Remarkably, exposure to morphine in cART-treated SIV-infected RMs led to a significant elevation of CD4+ Treg lymphocytes with a median of 5.22% versus 2.95% in the saline control group (p=0.004) (**Fig 4E**).

**Fig 4:**
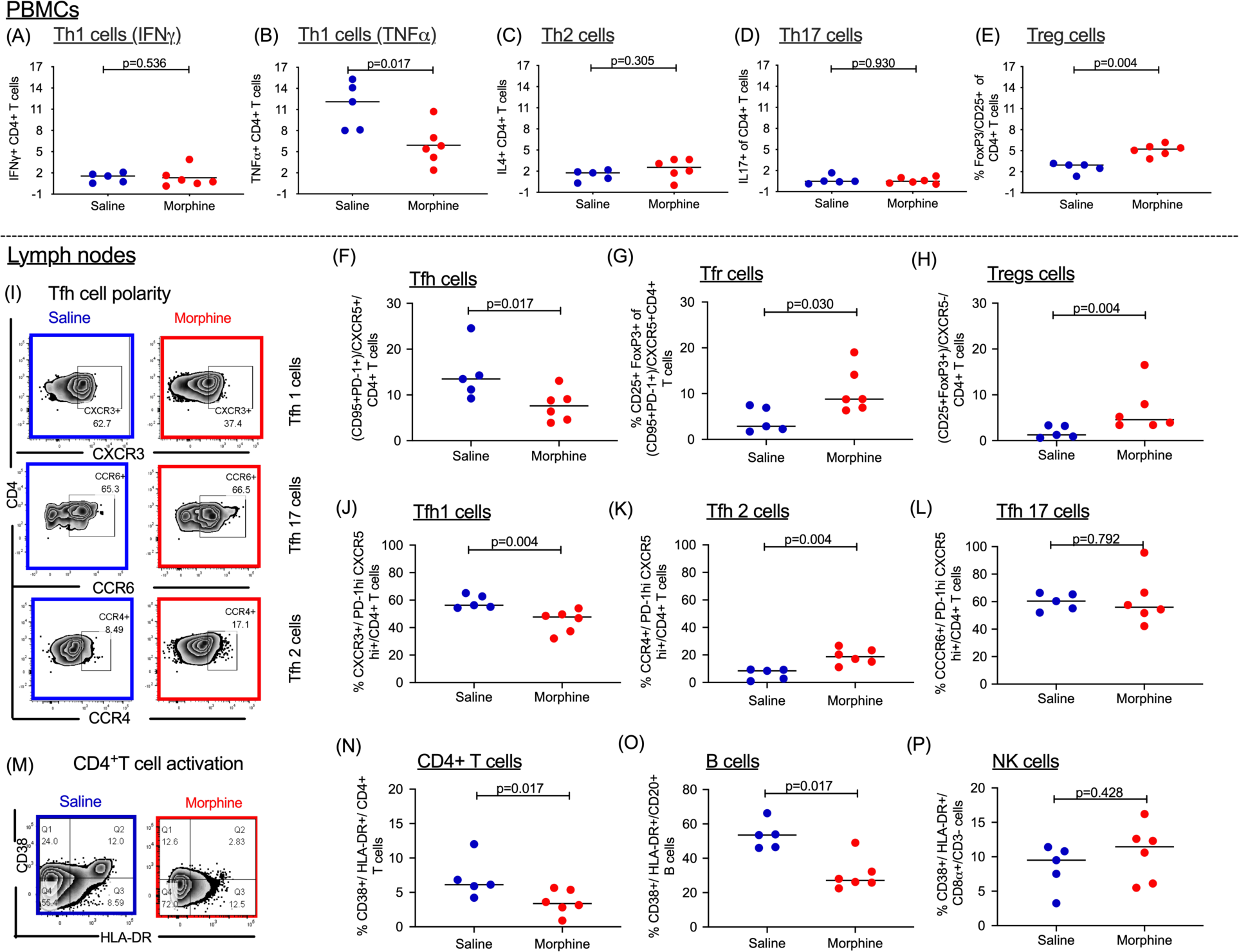
Immune dynamics in peripheral blood and lymph node compartments in morphine administered versus saline (control) groups of cART treated SIV mac 251 infected rhesus macaques. Following stimulation with PMA/ ionomycin, differences in the levels of: (A) PBMC CD4+ Th1 cells secreting IFNγ; (B) PBMC CD4+ Th1 cells secreting TNFα; (C) PBMC CD4+ Th2 cells secreting IL-4; (D) PBMC CD4+ Th17 cells secreting IL-17; (E) PBMC Tregs were also outlined as CD25+FoxP3+CD4+ T cells; Within the lymph nodes, the frequencies of (F) T follicular helper (Tfh) cells based on CD95+PD-1+ co-expression amongst CXCR5+ T cells were assessed. (G) Similarly, the levels of T follicular regulatory cells (Tfr) were evaluated based on CD25+ and FoxP3+ co-expression amongst the Tfh population. (H) Simultaneous evaluation of T regs (CD25+ and FoxP3+) within the CXCR5-CD4+ T cell population was also carried out. (I) Representation of Tfh polarity as delineated as Tfh1, Tfh 17 and Tfh 2 based on the surface expression of specific chemokine receptors. Collectively, (J) Tfh1 were based on CXCR3 (K) Tfh17 based on extent of CCR6 and (L)Tfh2 based on CCR4 expression amongst CXCR5 hi PD-1 hi Tfh cells. (M) Representation of CD4+ T cell activation in one experimental and one control sample. (N) Levels of CD4+ T cell (O) B cell and (P) NK cell activation as measured by the extent of CD38 and HLA-DR co-expression.

Within the LNs, numerous significant differences were found. Morphine administered cART treated RMs had lower frequencies of T follicular helper cells (Tfh) expressing PD-1, CXCR5 and CD95 with a median value of 7.61% versus 13.49% in saline control groups (p=0.017) (**Fig 4F**). On the other hand, in morphine administered group the frequency of regulatory Tfh (CD25+FoxP3+ Tfh) cells was elevated with a median of 8.78% versus 2.87% in saline control group (p=0.030) (**Fig 4G**). Similarly, there was an elevation of CD4+CXCR5-Treg (CD25+FoxP3+) cells in morphine administered group with a median of 4.57% versus 1.25% in saline control group (p=0.004) (**Fig 4H**). Evaluation of Tfh cell polarity was also carried out as shown by the representative sample in **Fig 4I**. There were differences in Tfh cell differentiation (Th1 vs Th2) in morphine administered group versus saline controls. In morphine group, while there was a reduction in CXCR3+Tfh cells (Th1) with a median of 47.70% versus 56.30% in saline control group (p=0.004) (**Fig 4J**), there was an elevation in CCR4+Tfh cells (Th2) with a median of 18.70% in the morphine administered group versus 8.49% in the saline control group (p=0.004) (**Fig 4K**). However, there were no differences in the frequency of CCR6+Tfh cells (Th17) with a median of 55.95% versus 60.40% in the morphine administered and saline control groups respectively (p=0.792) (**Fig 4L**). Finally, differences in immune activation across different cell subsets were evaluated. **Fig 4M** denotes representative sample for CD4+ T cell activation. For CD4+ T cell activation (denoted by CD4+/CD38+/ HLA-DR+), the morphine administered RMs had a significant lower frequency with a median of 3.37% versus 6.13% in saline controls (p=0.017) (**Fig 4N**). Similarly, for B cell activation (denoted by CD20+/CD38+/HLA-DR+), the morphine administered macaques had a significantly reduced frequency with a median of 27.10% versus 53.50% in saline controls (p=0.017) (**Fig 4O**). On the other hand, for NK cell activation (CD3-/CD8α/CD38+/HLA-DR+), no significant differences were observed in immune activation as shown by a median frequency of 11.45% in the morphine administered group versus 9.16% in the saline control group (p=0.428) respectively (**Fig 4P**).

### Quantification of SIV reservoirs in the CNS

To understand how chronic morphine administration impacted the seeding and persistence of viral reservoirs in the CNS, we analyzed CD11b+ myeloid cells (microglia and perivascular macrophages) from the brains of the cART-treated morphine administered and saline control SIV-infected monkeys, all of whom had complete viral suppression in the plasma and CSF (below 50 copies/ml). Immediately after necropsy CD11b+ cells were purified from brains. After purification, we obtained more than 90% cells positive for CD11b (**S6 Fig)**. The median of cell associated DNA load was 50.5 copies per million CD11b+ microglia/macrophages in the morphine administered group versus 0 in the saline control group (p=0.350) (**Fig 5A)**. The median cell-associated RNA load was 32.5 copies per million CD11b+ microglia/macrophages in morphine administered group versus 15 in the saline control group (p=0.710) (**Fig 5B)**. We next sought to estimate the size of latent replication-competent SIV reservoirs in CD11b+ microglia/macrophages using macrophage Quantitative Viral Outgrowth Assay (MØ-QVOA) as TILDA and IPDA is only validated to estimate viral reservoirs in CD4+ T cells. The number of CD11b+ microglia/macrophages used for the MØ-QVOA for each animal is described in **S2 Table**. The CD11b+ microglia/macrophages obtained from morphine administered macaques had a median infectious unit per million (IUPM) value of 0.89 versus 0.19 in saline control group (p=0.014) (**Fig 5C**), indicating presence of a significantly higher number of replication competent CD11b+ cells in the morphine group.

**Fig 5:**
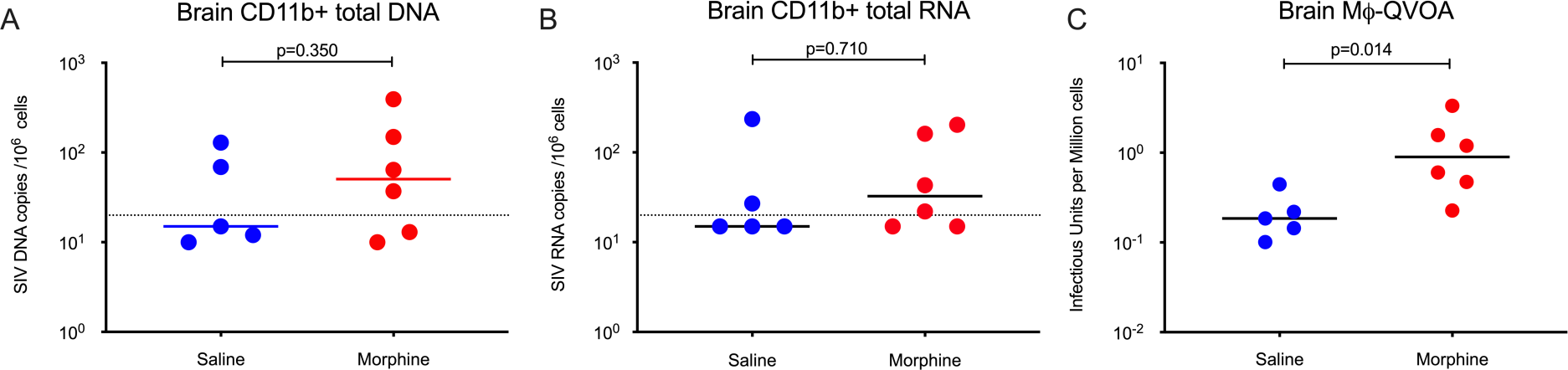
Size of SIV reservoirs in CD11b+ macrophages from brain of morphine administered versus control group of SIVmac251 infected ART suppressed *rhesus maca*ques. **(A)** SIV DNA copies per million CD11b+ macrophages isolated from brain of morphine administered and control rhesus macaques received saline; **(B)** SIV RNA copies per million CD11b+ macrophages isolated from brain of morphine administered and control rhesus macaques received saline. **(C)** Size of functional latent reservoirs estimated by macrophage quantitative viral outgrowth assay (MØ-QVOA) in CD11b+ positive macrophages purified from brain of morphine administered and control rhesus macaques received saline. Infectious unites per million cells (IUPM) were estimated using the IUPMStats v1.0 infection frequency calculator. All values below limit of detection are bringing to the limit of detection for better representation in the figure.

## Discussion

In order to achieve a functional cure of HIV-1, the primary goal is to identify all cell types and anatomical sanctuaries of viral reservoirs and to understand the molecular mechanism of their long-term persistence. Substantial work has been done to characterize and understand the underlying mechanism of latent reservoirs in circulating CD4+ T cells and in lymphoid tissues. Several line of evidence suggests that HIV enters the CNS within 3-7 days of infection and tissue resident macrophages and microglia, which likely serve as viral reservoirs and can be a source of rebound virus in case of therapeutic interruptions [33, 51-53]. The mechanism of persistence of HIV reservoirs in myeloid cells is still lacking [34]. Substance use disorders (SUD) and more specifically OUD is an important co-morbidity among PLWH [23]. Opioids have immune modulatory effects and mostly render macrophages more permissive to HIV infection [21, 24, 54, 55]. However, the impact of opioids on seeding and persistence of HIV reservoirs remains underscored, and this gap may pose a major roadblock in HIV cure research. To shed some light on this direction, in the present study we described an NHP model of HIV infection to understand how opioid use could modulate the size of viral reservoirs.

One result of the present study is that there was no significant difference in geometric mean of plasma and CSF viral loads between morphine administered versus control animals receiving saline in both untreated and ART suppressed groups. In contrast a previous study, Bokhari et al., reported that morphine administered rhesus macaques exhibited one log higher plasma and CSF viral load compared to controls. It is to be noted that in the Bokhari et al. study morphine was given four doses daily whereas we administered two daily doses. In addition, while we used SIVmac251, Bokhari et al., used a brain-derived stock of SIV (SIVmacR71/17E) [56]. In the Bokhari et al. study a high rate of rapid progression of disease (which is associated with higher viral loads) was found in the morphine treated group, similar findings were obtained by Kumar et al. using a mixture of different SIVs (SHIV(KU), SHIV(89.6)P and SIV/17E-Fr) [57]. Furthermore, as the rhesus macaques used in different studies are outbreed from multiple colonies their genetic background may vary, and subsequently their response to morphine administration and SIV disease pathogenesis. These above differences could have been contributed the differences of plasma and CSF viral loads observed in these studies.

Among the untreated RMs, we did not find any profound difference in cell-associated DNA and RNA loads at different tissue compartments between morphine administered and the saline control group, however, there was trend of lowered viral loads in morphine treated animals. On the other hand, in cART suppressed RMs, we observed a reduction in cell-associated DNA and RNA load in CD4+ T cells obtained from peripheral blood and LNs as well as in tissue samples collected from rectal mucosa in morphine administered macaques compared to saline controls. This finding is substantiated by IPDA and TILDA performed on CD4+ T cells purified from PBMC and LNs, where the number of intact SIV genome per million CD4+ T cells is less and the size of inducible SIV reservoirs in CD4+ T cells is lowered significantly in morphine administered RMs compared to saline controls, respectively. These findings indicate that chronic morphine administration in combination with ART has a positive impact in lowering SIV reservoirs in lymphoid tissues.

CD4+ T cells are major source of long-term latent reservoirs of HIV in lymphoid tissues like LNs, GALT and in peripheral blood among others [58]. In LNs HIV reservoirs seeded during acute infection, associated with virus production and storage of viral particles in immune complexes, and Tfh cells constitute the major part of LN viral reservoirs [59-61]. We observed depletions of Tfh cells in LNs of morphine administered macaques that supports earlier reports that chronic morphine abuse depletes the volume and total number of lymphoid cells from LNs [62, 63]. Brown et al., reported that morphine induce an immune-suppressive effect in LNs and peripheral blood of African green monkeys and pigtailed macaques with a substantial decrease in abundance of several metabolic proteins involved in energy metabolism pathways accompanied by significant decreases in activated CD4^+^/ CD8^+^ T cells [64]. Therefore, we speculate, in our experimental model chronic morphine administration may dampen the T-cell activation and suppress their metabolic activity, cause a reduction in seeding of SIV CD4+ T cells during acute stage of infection before initiation of cART, which may result in smaller size of persistent latent SIV reservoirs in CD4+ T cells of morphine administered animals compared to control macaques received saline.

Next, we observed that morphine administration resulted in the expansion of regulatory T cells in peripheral blood as well as in LNs, that have been previously reported to suppress T-cell activation [65]. In LNs, we also observed lower level of activation of CD4+ T cells and B cells in morphine administered animals compered to saline controls. Which supports the proposed mechanism of morphine mediated reduction on SIV reservoirs within the lymphoid tissue compartments. Furthermore, recently Lee et al., reported that in chronic HIV patients with suppressed plasma viral load, intact replication competent proviruses persist in Th1-polarized memory CD4+ T cells [47]. Interestingly, chronic morphine exposure leads to a phenotypic switch of Th1 to Th2 in CD4+ T cells [37]. We also observed a lower frequency of IFNγ and TNFα secreting Th1 polarized cells in peripheral blood of morphine administered RMs. Similarly, in LNs of morphine administered animals we observed a reduction in frequency of CXCR3+Th1 like Tfh cells and expansion of CCR4+Th2 like Tfh cells and it was reported earlier that Th1 like Tfh cells enters LN germinal center during acute stage of HIV/SIV infection, express high level of CCR5 and constitute major part of viral reservoirs in LNs [66-68]. In combination of the above facts may leads to depletion of functional SIV reservoirs in CD4+ T cells of morphine administered ART suppressed rhesus macaques (**Fig 6A**).

**Fig 6:**
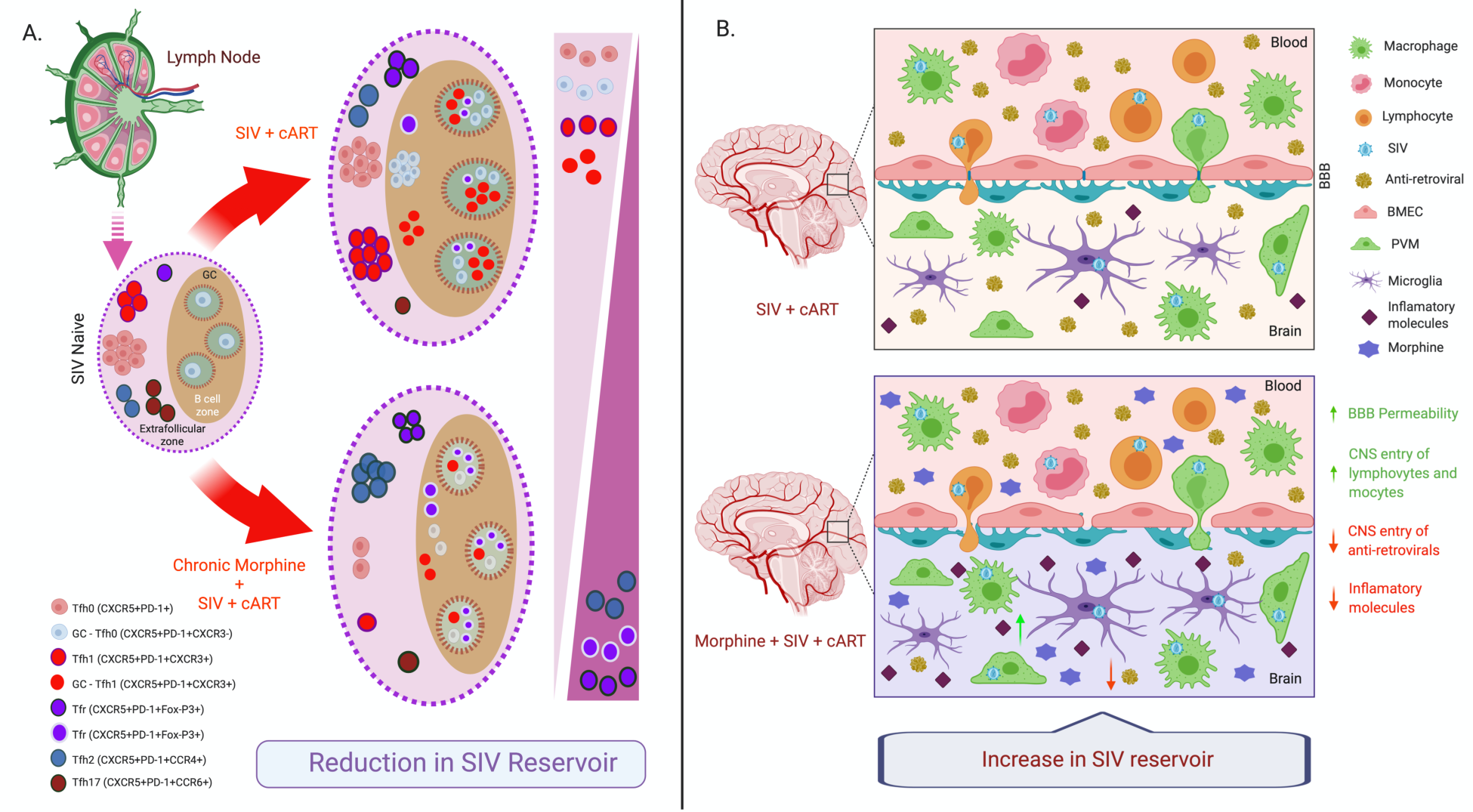
Proposed mechanism of morphine mediated differential regulation of SIV reservoirs in lymphoid tissue versus myeloid cells in CNS. **(A)** Chronic morphine exposure leads to a reduction of Tfh cells in lymph nodes (LNs) which serves as the major HIV/SIV reservoirs in LNs. This may be responsible for observing reduction in SIV reservoirs in our morphine dependent, SIVmac251 infected cART treated *rhesus macaque* model. **(B)** Chronic morphine exposure increases the permeability of blood brain barrier (BBB), which increases the trafficking of lymphocytes and monocytes to the CNS. Again, opioids lower the CNS penetration of several anti-retroviral drugs to facilitate basal level of ongoing HIV replication in brain. This may be responsible for higher size of functional SIV reservoirs in our morphine dependent SIVmac251 infected cART treated *rhesus macaque* model.

Contrary to the CD4+ T cells we observed an opposite effect of morphine in modulation of the viral reservoir in CNS. The median size of replication competent SIV reservoirs in CD11b+ microglia/macrophages from brain is significantly higher in morphine administered RMs compared to control animals receiving saline. There are several factors that may be responsible for the morphine-mediated modulation of CNS SIV reservoirs. It has been reported that morphine upregulates the expression of CCR5 coreceptor in macrophages, which, in turn, could increase the viral permissiveness of these cells [21, 55, 69]. Morphine is known to also lower the CNS penetration of several antiretrovirals, which could result in ongoing viral replication in the CNS and responsible for larger reservoir [70]. Again, it has been shown that morphine alone or in combination with HIV TAT increases the permeability of blood brain barrier (BBB) and increases trans-endothelial migration of leucocytes by activation of proinflammatory cytokines, intracellular Ca_2_+ release and activation of myosin light chain kinase that down regulate tight junction proteins and decreased transendothelial electric resistance [26, 71, 72]. It also increases the release of CCL2, CCL5 and IL-6 by astrocytes and upregulates expression of ICAM and VCAM on brain vascular endothelial cells that promotes trafficking of HIV infected peripheral leucocytes in CNS [73]. Morphine is also shown to down regulates the expression of anti-HIV microRNAs and impair the function of anti-HIV restriction factors in monocytes [74, 75]. Taken together, this could lead to increased size of the viral reservoirs in the CNS of morphine administered group in comparison to the controls received saline. We observed a similar analogy with our finding of differential modulation of SIV reservoirs by morphine in T cells from PBMC/LNs versus microglia/macrophages in CNS in some previous reports where phenelzine, a monoamine oxidase (MAO) inhibitor, suppresses reactivation of HIV in T cells [76] and reactivates HIV in human microglia cell lines [77]. To the best of our knowledge this is the first report of opioid mediated differential modulation of HIV reservoirs using macaque models of HIV infection (**Fig 6B)**.

The major caveat of the current study is the number of RMs included in each group due to the prohibitive cost of conducting long-term studies with multiple daily injections. Other issues include the experimental design where animals were initially ramped-up with morphine, then infected and then treated with cART. In a patient cohort, this may not be ideal scenario, as most of the people suffering from substance use disorders take multiple forms of drugs. For some of the PLWH, opioids are prescribed to treat chronic pain, and subsequently they become addicted to the substance. For this group of population how opioids use modulates viral reservoirs dynamics needs to be understood and these will be interest of our future studies.

In conclusion, for the first time we describe a morphine dependent SIVmac251 infected RM model to study the impact of opioid use disorder (OUD) on HIV reservoirs. Our results suggest that morphine differentially modulate SIV reservoirs in LNs and peripheral blood versus SIV reservoirs in myeloid lineage of cells within the CNS. Chronic morphine administration reduces the size of viral reservoirs in CD4+ T cells, whereas increases the size of SIV reservoirs in brain resident CD11b+ macrophages. We propose that, these pre-clinical models will serve as a tool to discover the molecular mechanism of opioid mediated differential regulation among PLWH and suffering from OUD.

## Methods

### Reagents and cell lines

Antiretroviral drugs tenofovir alafenamide (TFV) and emtricitabine (FTC) were obtained from Gilead Sciences, Foster City, CA, USA, while dolutegravir (DTG) was procured from ViiV Healthcare, Research Triangle Park, NC, USA as MTA with Dr. Byrareddy and UNMC. All other molecular biology grade fine chemicals used in the study were purchased from Sigma-Aldrich, St. Louis, MO, USA if otherwise mentioned specifically. All primers and probes used in this study was custom synthesized from Integrated DNA Technologies, Inc (Coralville, Iowa; USA). CEMx174 is a hybrid human lymphoid cell line generated from human B721.174 and T-CEM cell lines, that was used for expansion of virus in quantitative viral outgrowth assays due to its enhanced susceptibility to SIV infection [78]. This cell line was provided by Dr. J. Hoxie (University of Pennsylvania, Philadelphia). CEMx-174 cells were maintained in complete RPMI (RPMI-1640 (Gibco; Cat# 21870076), 10% heat-inactivated FBS (Gibco; Cat#10437-028), 2 mM L-Glutamine (Gibco; Cat#35050079) and 100 U/ml penicillin and 100 ug/ml streptomycin (Gibco; Cat#15140-122) at 37°C and 5% CO_2_.

### Animals and Ethical statement

A total nineteen Indian origin, outbred, pathogen free RMs (*Macaca mulatta*, mean age: 4.67 years range: 4.1 to 7.0 years) were used in this study (**Table 1**). The macaques were housed in compliance with the regulations under the Animal Welfare Act, the Guide for the Care and Use of Laboratory Animals in the nonhuman primate facilities at the Department of Comparative Medicine, University of Nebraska Medical Center (UNMC), Omaha, Nebraska, USA. Animals were maintained in a temperature controlled (72°F) indoor climate with 12-hour light/dark cycle. The monkeys were observed twice daily for development of distress or disease by the animal care staffs and veterinary personnel. The animals were daily fed monkey diet (Purina) supplemented with fresh fruit or vegetables and water ad libitum.

**Table 1:**
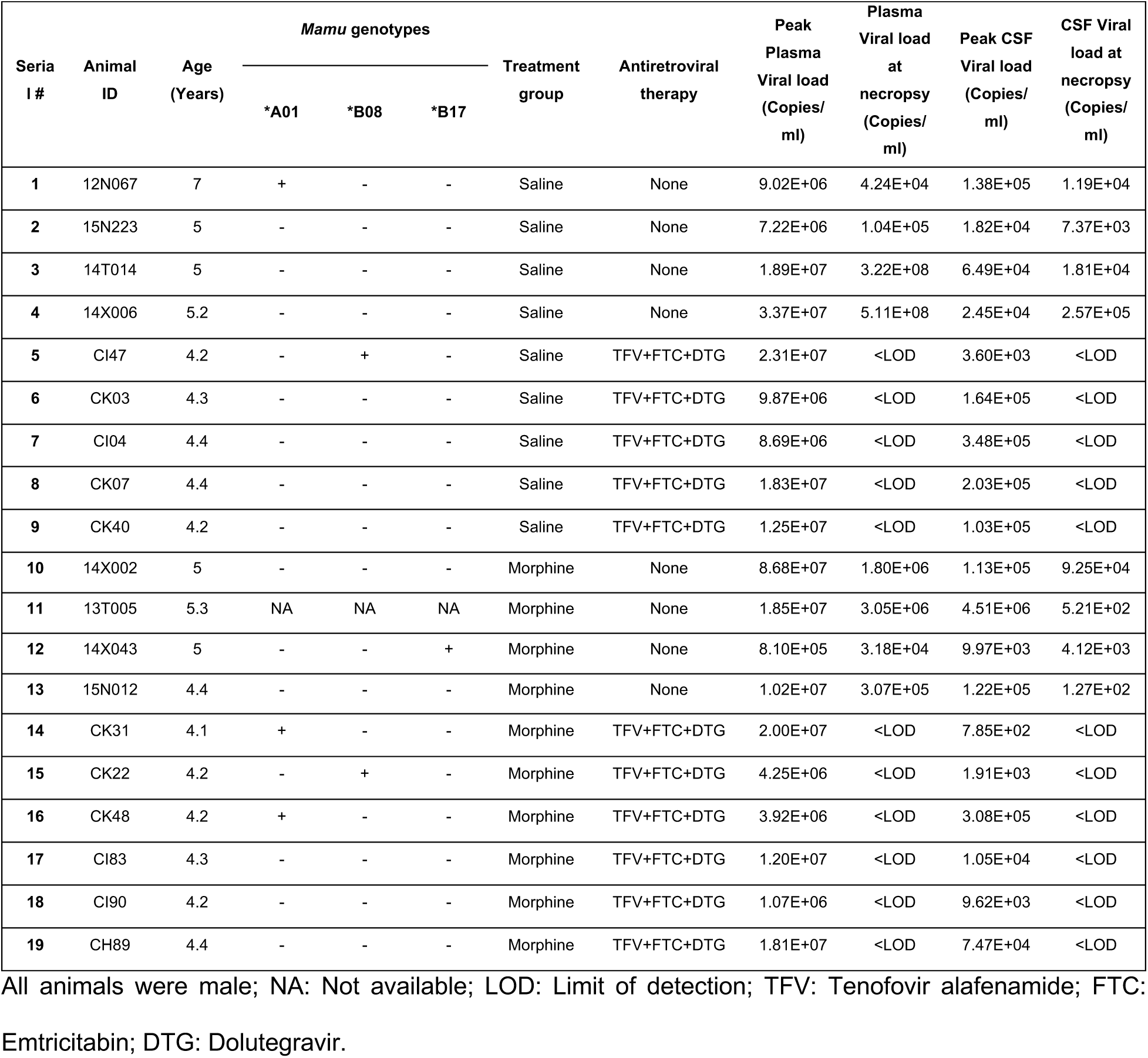
Description of rhesus macaques used in the study

| Serial # | Animal ID | Age (Years) | Mamu genotypes |  |  | Treatment group | Antiretroviral therapy | Peak Plasma Viral load (Copies/ml) | Plasma Viral load at necropsy (Copies/ml) | Peak CSF Viral load (Copies/ml) | CSF Viral load at necropsy (Copies/ml) |
| --- | --- | --- | --- | --- | --- | --- | --- | --- | --- | --- | --- |
|  |  |  | *A01 | *B08 | *B17 |  |  |  |  |  |  |
| 1 | 12N067 | 7 | + | - | - | Saline | None | 9.02E+06 | 4.24E+04 | 1.38E+05 | 1.19E+04 |
| 2 | 15N223 | 5 | - | - | - | Saline | None | 7.22E+06 | 1.04E+05 | 1.82E+04 | 7.37E+03 |
| 3 | 14T014 | 5 | - | - | - | Saline | None | 1.89E+07 | 3.22E+08 | 6.49E+04 | 1.81E+04 |
| 4 | 14X006 | 5.2 | - | - | - | Saline | None | 3.37E+07 | 5.11E+08 | 2.45E+04 | 2.57E+05 |
| 5 | CI47 | 4.2 | - | + | - | Saline | TFV+FTC+DTG | 2.31E+07 | <LOD | 3.60E+03 | <LOD |
| 6 | CK03 | 4.3 | - | - | - | Saline | TFV+FTC+DTG | 9.87E+06 | <LOD | 1.64E+05 | <LOD |
| 7 | CI04 | 4.4 | - | - | - | Saline | TFV+FTC+DTG | 8.69E+06 | <LOD | 3.48E+05 | <LOD |
| 8 | CK07 | 4.4 | - | - | - | Saline | TFV+FTC+DTG | 1.83E+07 | <LOD | 2.03E+05 | <LOD |
| 9 | CK40 | 4.2 | - | - | - | Saline | TFV+FTC+DTG | 1.25E+07 | <LOD | 1.03E+05 | <LOD |
| 10 | 14X002 | 5 | - | - | - | Morphine | None | 8.68E+07 | 1.80E+06 | 1.13E+05 | 9.25E+04 |
| 11 | 13T005 | 5.3 | NA | NA | NA | Morphine | None | 1.85E+07 | 3.05E+06 | 4.51E+06 | 5.21E+02 |
| 12 | 14X043 | 5 | - | - | + | Morphine | None | 8.10E+05 | 3.18E+04 | 9.97E+03 | 4.12E+03 |
| 13 | 15N012 | 4.4 | - | - | - | Morphine | None | 1.02E+07 | 3.07E+05 | 1.22E+05 | 1.27E+02 |
| 14 | CK31 | 4.1 | + | - | - | Morphine | TFV+FTC+DTG | 2.00E+07 | <LOD | 7.85E+02 | <LOD |
| 15 | CK22 | 4.2 | - | + | - | Morphine | TFV+FTC+DTG | 4.25E+06 | <LOD | 1.91E+03 | <LOD |
| 16 | CK48 | 4.2 | + | - | - | Morphine | TFV+FTC+DTG | 3.92E+06 | <LOD | 3.08E+05 | <LOD |
| 17 | CI83 | 4.3 | - | - | - | Morphine | TFV+FTC+DTG | 1.20E+07 | <LOD | 1.05E+04 | <LOD |
| 18 | CI90 | 4.2 | - | - | - | Morphine | TFV+FTC+DTG | 1.07E+06 | <LOD | 9.62E+03 | <LOD |
| 19 | CH89 | 4.4 | - | - | - | Morphine | TFV+FTC+DTG | 1.81E+07 | <LOD | 7.47E+04 | <LOD |

All animals were male; NA: Not available; LOD: Limit of detection; TFV: Tenofovir alafenamide; FTC:
Emtricitabin; DTG: Dolutegravir.

At the end of the study, all RMs were humanely euthanized using a high plane of anesthesia with ketamine/zylazine and then opening the thoracic cavity and perfused/exsanguinated with sterile PBS, in accordance with the guidelines of the American Veterinary Medical Association. This study was reviewed and approved by UNMC Institutional Animal Care and Use Committee (IACUC) and the Institutional biosafety Committee (IBC) under protocol number ‘16-073-07-FC’ titled “The effect of cART and drug of abuse on the establishment of CNS viral reservoirs” and ‘15-113-01-FC’ titled “The combinatorial effects of Opiates and promoter-variant strains of HIV-1 subtype C on neuropathogenesis and latency”. UNMC has been accredited by the Association for Assessment and Accreditation of Laboratory Animal Care International.

### Study Design

The overall design of the study is illustrated in a schematic form in **S1 Fig**. These experiments were designed to investigate the effect of chronic morphine administration on the establishment of viral reservoir in SIV-infected RMs in different anatomical compartments of the body. The study included 19 juvenile RMs. The RMs were randomly divided into two groups. One group with ten animals were ramped-up over two weeks to a final 5 mg/kg intramuscular injection of morphine twice daily, which was maintained for seven weeks and the other group (n=9) received similar dose of normal saline (control group). At this point all the macaques were intravenously inoculated with 200 TCID_50_ of SIVmac251 (viral stock was obtained from Dr. Mahesh Mohan from Tulane National Primate Research Center), while the administration of morphine/saline continued till the end of the study. After five weeks post-inoculation, daily ART was initiated in six macaques from morphine group and five macaques in the control group and continued till the end of the study. ART regimen consisted of two reverse transcriptase inhibitors (FTC: 40 mg/ml and TFV: 20 mg/ml) and one integrase inhibitor (DTG: 2.5 mg/ml). All the drugs were dissolved in a vehicle made up of 15% Kleptose HPB (Roquette, parenteral grade) (w/w) in 0.1N NaOH. The anti-retroviral drugs were administered subcutaneously once daily at 1 ml/kg body weight. Peripheral blood from the femoral vein, CSF by direct puncture of the cisterna magna or by lumbar puncture, LNs and colorectal mucosa biopsies were collected longitudinally at different time points of the study after anesthetizing the monkeys with ketamine-HCl (5-20 mg/kg) or telazol (3-5 mg/kg) to monitor SIV viral loads and a series of immunologic and virologic parameters as described in experimental schema.

### SIV plasma and CSF viral load quantification

SIV RNA concentration in the plasma and CSF samples were measured by quantitative reverse transcription-PCR (qRT-PCR) as previously described [79]. In brief, plasma was separated from blood samples collected in K2-EDTA vacutainer tubes (Becton Dickinson, San Diego, California, USA) within 4 hours of collection. RNA was extracted from 140 µl of plasma and CSF samples using a QIAamp Viral RNA mini kit according to manufacturer’s instructions (QIAGEN Germantown, MD, USA; Cat# 52906). SIV gag RNA was quantified by qRT-PCR using the TaqMan RNA-to-Ct 1-Step Kit (Thermo Fisher Scientific, MA; Cat# 4392938) and Applied Biosystems QuantStudio 3 Real-Time PCR System (Applied Biosystems, Waltham, MA, USA). Primers and probes used for SIV gag RNA quantification were as follows: SIVGAGF: 5’ – GTCTGCGTCATCTGGTGCATTC – 3’; SIVGAGR: 5’ – CACTAGGTGTCTCTGCACTATCTGTTTTG – 3’ and SIVP: 5’-/6-FAM/ CTTCCTCAG /ZEN/ TGTGTTTCACTTTCTCTTCTGCG /3IABkFQ/ - 3’.

### Purification of CD4+ T cells

Peripheral blood mononuclear cells (PBMCs) and LN cells were enriched for CD4+ T cells using the EasySep NHP CD4+ T cell isolation kit from STEMCELL Technologies Canada Inc (Cat#: 19582) as per manufacturers instruction. Briefly, frozen cells were thawed at 37°C, washed with complete RPMI (composition described above), centrifuged for 6 min at 1200rpm and resuspended in 1 ml of recommended medium (Phosphate Buffered Saline (PBS) containing 2% Fetal Bovine Serum (FBS) and 1 mM EDTA). The cell suspension was transferred to a 12×75 mm polystyrene tube, 50 µl of EasySep Negative Selection cocktail was added, mixed well and the cell suspension were incubated at room temperature (RT) for 10 minutes. Then 100 µl of EasySep Magnetic Particles added and the cells were incubated at RT for 5 minutes. After incubation, 2 ml of recommended medium was added to the solution, then the tubes were placed in an EasySep magnet for 5 min. The negatively selected enriched CD4+ T cells were collected for downstream applications.

### Isolation of total myeloid-enriched brain cells

Myeloid-enriched brain isolation used a modification of the procedure described by Marcondes et al., [80]. In brief, the brain was sectioned and meninges removed in Hank’s balanced salt solution (HBSS; Invitrogen, Carlsbad, CA). The cleaned-tissue was homogenized with a dounce homogenizer and washed twice with HBSS. Brain homogenate was digested on a rotating platform at 37° C for 30 minutes in HBSS with 28 U/ml DNase1 and 8 U/ml papain (Sigma, St. Louis, MO). The samples were triturated after 15 and 30 minutes digestion. Enzymes were inactivated with addiion of 3 % FBS. Sample was washed twice with HBSS and then centrifuged for 15 minutes at 1800 RPM at 4° C through 25% Percoll (GE HealthCare, Pittsburg, PA). The resulting fatty upper and fluid middle layers were removed from the pellet. The pellet was resuspended in 10ml of ice-cold HBSS and filtered through a 40 µm screen. Brain isolates were counted on a hemocytometer and viability was measured by Trypan blue exclusion.

### Enrichment of CD11b myeloid cells

Cell isolates were washed in PBS, reconsituted in MACS buffer with 0.1% BSA (Miltenyi, Gladbach, Germany). Cells were counted and volume was adjusted for staining with non-human primate CD11b microbeads (Miltenyi, Gladbach, Germany). Forty million cells were reconstituted in 160 µl of MACS buffer and reacted with 80 µl of CD11b microbeads at 4°C for 15 minutes. After incubation, cells were washed with MACS buffer with 0.1% BSA, reconstituted into 500 µl of MACS buffer and loaded onto MACS Separator LD columns. Both the negative flow through and the positve CD11b cells were collected, cells were counted on Coulter Counter Z1. Isolates were used for downstream processing and analyzed for epitope expression using FACS.

### Purification of alveolar macrophages from Bronchoalveolar lavage (BAL)

At necropsy the lung was lavaged the with sterile saline and the BAL collected. Cells were collected through centrifugation of the BAL at 250 g for 10 min at 4°C. The cell pellets were washed twice with PBS containing 2% FBS and 2 mM EDTA. Cells were resuspended in complete RPMI (composition described above) and used for downstream applications.

### Isolation DNA/RNA from CD4+/CD11b+ cells

Enriched CD4+ T cells and CD11b+ macrophages were pelleted by centrifugation for 6 minutes at 1200 rpm and stored at -80°C until time of processing. To extract the DNA/RNA from the cells, first the pellets were thawed on ice and thoroughly loosened by flicking the tube. Then, 600 µl Buffer RLT Plus with 2-Mercaptoethanol (β-ME) at 10 µl/1 ml concentration was added and thoroughly mixed by vortexing. Following lysis, the mixture was transferred to a QIAshredder (QIAGEN, Cat#79656) and the AllPrep DNA/RNA Mini Kit (Qiagen, Cat#80204) was used for DNA/RNA isolation. The remaining protocol followed was per the kit’s instruction. RNA was eluted in 30 µl of RNase free water after performing column DNase digestion while DNA was eluted in 50 µl of Elution Buffer provided in the kit. DNA/RNA concentrations were measured using SimpliNano spectrophotometer (GE Healthcare Bio-Sciences Corp., Piscataway, NJ, USA) and stored at -80°C deep freezer for future use.

### Isolation DNA/RNA from frozen tissues

For DNA/RNA isolation from tissues, the AllPrep DNA/RNA Mini Kit (QIAGEN Germantown, MD, USA; Cat#80204) was used. Approximately 30 mg of frozen tissue was placed into a 2 ml flat bottom centrifuge tube on dry ice with a 5 mm stainless steel bead (QIAGEN Germantown, MD, USA; Cat#69989) at the bottom. Then, 600 µl buffer RLT plus with 2-Mercaptoethanol (βME) at 10 µl βME/ml of RLT+ was added before placing the tube into a TissueLyser LT (QIAGEN Germantown, MD, USA; Cat#69980) set at 50 oscillations/second for 5 minutes. The lysate was checked for homogeneity and transferred to a QIAshredder (QIAGEN Germantown, MD, USA; Cat#79656). DNA/RNA was then isolated per the kit’s instruction. RNA was eluted in 30 µl of RNase free water after performing column DNase digestion while DNA was eluted in 50 µl of buffer EB. DNA/RNA concentrations were measured using SimpliNano spectrophotometer (GE Healthcare Bio-Sciences Corp., Piscataway, NJ, USA) and stored at -80°C deep freezer for future use.

### SIV cell associated RNA/DNA quantification

Total cell associated SIV RNA was quantified using RT-ddPCR using 100 nanogram of RNA, 1-Step RT-ddPCR Advanced Kit for Probes (Bio-Rad; Cat#1864022) and same set of primers and probe used for SIV plasma viral load quantification. ddPCR was carried out on Bio-Rad QX200 AutoDG digital droplet PCR system. In brief, 22 µl of reaction mix was used for droplet generation using QX200™ Droplet Generator, the ddPCR plate having the emulsified samples was heat sealed with a foil (Bio-Rad; Cat#181–4040) and amplified in a C1000 Touch thermal cycler (Bio-Rad, CA, USA). After thermal cycling, ddPCR plates were transferred to the QX200 droplet reader (Bio-Rad, CA, USA) for droplet count and fluorescence measurement. Positive droplets with amplified products were separated from negative droplets without target amplicon by applying a fluorescence amplitude threshold and the absolute quantity of RNA per sample (copies/µl) was determined using QuantaSoft software. For quantification of cell associated SIV DNA the same methodology was used as described above without using the reverse transcription step and used 2X ddPCR Supermix for Probes (no dUTP, Bio-rad; CA, USA; Cat#1863024) instead of 1-Step RT-ddPCR Advanced Kit.

### Tat/rev Induced Limiting Dilution Assay (TILDA)

Tat/rev Induced Limiting Dilution Assay (TILDA) was performed as described by Frank et al., [81]. In brief, enriched CD4+ T cells from frozen PBMCs or LN total cells were suspended in complete RPMI at 2 million cells/ml concentration, and then rested at 37°C, 5% CO2 for a minimum of 2 hours. After resting, cells were divided in two group, one was stimulated with 100 ng/ml PMA and 1ug/ml ionomycin, and the other group left untreated as control. After 16 hours, the cells were counted and then centrifuged at 2000 rpm for 10 minutes to stop the reaction. Cells were then serially diluted in complete RPMI to the following: 18,000, 9,000, 3,000, and 1,000 cells/ul. A 96-well PCR plate was divided column-wise into thirds for the three treatments, and then into quarters for the four dilutions set in 8 replicates. In each well of the plate, 5 µl of Master Mix was added and then 1 µl of the appropriate dilution for the corresponding treatment was added to each well. The first PCR reaction mix contained 0.2 µl of Super Script III RT/Platinum Taq Mix (Invitrogen; Cat# 2010520), 0.1 µl of Superase Rnase Inhibitor (Invitrogen; Cat#00749138), 2.25 µl of Nuclease Free H2O (Ambion, Cat#AM9937), 2.2 µl of tris-EDTA (TE) Buffer, 0.125 µl of 10 µM SIVF (5’ - CACGAAAGAGAAGAAGAACTCCG – 3’, IDT) and 0.125 ul 10 µM SIVR (5’ - TCTTTGCCTTCTCTGGTTGG – 3’, IDT). To each well of the plate, 5 µl of reaction mix were added. Pre-amplification settings included reverse transcription for 15 min at 50°C, followed by denaturation for 2 min at 95°C, then 24 cycles of amplification at 95°C for 14 seconds and 60°C for 4 minutes. Pre-amplification was done in a LifePro Thermal Cycler (Bioer Technology). After pre-amplification, 9 µl of qPCR reaction mix containing 5 ul LightCycler 480 Probes Master (Roche, Cat#04707494001), 3.4 ul of Nuclease Free H2O (Ambion, AM9937), 0.2 µl of 20 µM Nested FW primer (5’ - AGGCTAAGGCTAATACATCTTCTG – 3’, IDT), 0.2 µl of 20 µM SIVR (5’ - TCTTTGCCTTCTCTGGTTGG – 3’, IDT), and 0.2 µl of 5 µM PROBE (5′-/56-FAM/AAACCCATA/ZEN/TCCAACAGGACCCGG/3IABkFQ/-3′) details were added to each well of a new 96-well PCR plate. Then, 1 µl of PCR product from pre-amplification was added to the corresponding well in the new plate. Real-time PCR was performed in QuantStudio 3 (Applied Biosystems, Waltham, MA, USA) with the following settings: Pre-incubation at 95°C for 10 min, followed by 45 cycles at 95°C for 10 sec, 60°C for 30 sec, and 72°C for 1 sec, then finally a cooling period at 40°C for 30 sec.

### Intact proviral DNA assay (IPDA)

Intact proviral DNA assay was performed as described by Bender et al. [46], including thermal cycling conditions, primers and probes used in this assay. In brief, 250-1000 ng of DNA was mixed with 10 µl of 2X ddPCR Supermix for Probes (no dUTP, Bio-rad; Cat#1863024), 600 nM of primers and 200 nM of probes. ddPCR was carried out on Bio-Rad QX200 AutoDG digital droplet PCR system as described above. To quantify the input cell numbers, a parallel ddPCR reaction for rhesus macaque RPP30 gene was carried out.

### Macrophage Quantitative Viral Outgrowth Assay (MØ-QVOA)

In this study, a modified version of macrophage (MØ-QVOA) was performed on CD11b+ macrophages isolated from brain, lung, and liver as described by Avalos et al., [82]. In brief, cells were purified using a CD11b+ isolation kit per the manufacturer’s recommended protocol (Miltenyi Biotec, Auburn, CA; cat# 130-091-100). Prior to setting the culture, plates were coated with poly-L-lysine solution (Sigma, NC9778696) for 30 min and then washed twice with PBS. Purified cells were plated in triplicate and a 10-fold serial dilution in the presence of 10 µM zidovudine (Sigma), 25 nM DRV (Janssen), and 5 nM RTV (Merck). BrMφ media (DMEM (Gibco; Cat#12491-015), 5% heat-inactivated FBS (Gibco; Cat#10437-028), 5% IS giant cell tumor conditioned media (Irvine Scientific; Cat#91006), 100 U/ml penicillin and 100 ug/ml strep (Gibco; Cat# 15140-122), 70 µg/ml gentamycin (Sigma Aldrich; Cat#G1397-10ML), 2 mM L-glutamine (Gibco; Cat# 35050079), 3 mM sodium pyruvate (Gibco; Cat#11360070) and 10 mM HEPES buffer (Gibco; Cat#35050079)) was used for brain cells and complete RPMI for lung and liver. The culture was then placed in a 37° C, 5% CO2 incubator for 72 hours to permit proper cell adherence. After the 3-day incubation, the antiretroviral-containing medium was removed and cells were washed with PBS (1X) and then replenished with BrMφ/complete RPMI media containing 10 ng/ml tumor necrosis factor alpha (TNF-α; ProSpec; Cat#CYT-114), 1 µg/ml Pam3csk4 (Sigma Aldrich; Cat# 506350), and 1 µg/ml prostaglandin (Sigma Aldrich; Cat#538904). Additionally, approximately 10^5^ CEMX-174 feeder cells were added to each well. Supernatants were collected on days 5, 7, 10, and 14 post-culture activation and replenished with activating agent’s TNF-α, Pam3csk4, and Prostaglandin containing media. RNA was extracted from the supernatants using the QIAamp Viral RNA Mini Kit (QIAGEN, Cat#52906). Prior to extraction, the supernatants were centrifuged at 2000 rpm for 5 minutes, and then 140 µl was aliquoted and used per the kit’s instruction to isolate RNA. The RNA was eluted in 50 µl of AVE buffer. Viral RNA was quantified in culture supernatant using RT-ddPCR as described above. The frequency of replication competent latent reservoir cells was estimated using the IUPMStats v1.0 infection frequency calculator [83].

### Flow cytometry evaluation of changes in T cell polarization in peripheral blood and lymph nodes

The details of all fluorochrome-conjugated monoclonal antibodies used in the flow cytometry experiments is listed in **S1 Table**. the Single cell suspensions of PBMC and lymph nodes were prepared from samples collected from cART treated rhesus macaques (n=11) during necropsy. 2-4 million of PBMCs were stimulated with PMA/ionomycin at a final concentration of 50 ng/ml and 500 ng/ml respectively in complete RPMI-1640 (composition described above). An unstimulated condition was included as an experimental control. After 2 hours of incubation at 37°C in a humidified 5% CO2 incubator, 1 μl of GolgiPlug™ (BD Biosciences, USA; cat#555029) containing brefeldin A was added to each well together with 1X of monensin (Biolegend, San Diego, CA; cat#420701) and incubated for an additional 4 hours. After that 1/1000 dilution of zombie aqua amine reactive dye (Biolegend, San Diego, CA; cat# 423101) was added to the cell suspensions and incubated for 30 minutes in the dark, cells were washed and Fc blockade carried out using polyclonal anti-human Fc Receptor binding inhibitor (eBioscienceTM, USA; Cat#16-9161-71). Surface staining was performed using anti-CD45, anti-CD8, anti-CD4 and anti-NKG2A for panel 1 and using anti-CD25 and anti-CD127 for panel 2 (Treg panel) and cells were incubated for 1 hour at room temperature in dark. Thereafter for panel 1, cells were fixed using 2% paraformaldehyde (PFA) solution for 30 minutes and permeabilization was performed using a 1X BD perm/wash solution (BD Biosciences, USA; cat#554723) for 15 minutes at 4°C. Then, anti-CD3, anti-IFNγ, anti-TNFα and anti-IL-17 antibodies were added, and cells were incubated at 4°C for 15 minutes, washed and resuspended in PBS. For the Treg panel, fixation and permeabilization was performed using a 1X solution of Foxp3/Transcription Factor Fix/Perm Concentrate (4X) (Tonbo Biosciences, San Diego, CA; Cat#TNB-1020-L050) that was diluted using the 1X Foxp3/Transcription Factor Fix/Perm Diluent (Tonbo Biosciences, San Diego, CA; Cat#TNB-1022-L160). Following this, anti-CD3, anti-IL-4 and anti-FOXP3 antibody were added and cells were incubated at 4°C for 15 minutes, washed and resuspended in PBS.

For LN cell suspensions, 2-4 million cells were taken, and zombie UV viability dye was added, and Fc receptor blockade performed as described above. Then, in the T cell polarization panel, anti-CXCR5, anti-CXCR3, anti-CCR4 and anti-CCR6 antibodies were added and in the Treg panel anti-CXCR5 antibody was added. After that cells in both the panel was incubated at 370C for 30 minutes. Following this, ani-CD45, anti-CD3, anti-CD4, anti-CD8, anti-CD20, anti-CD95, anti-CD38, anti-PD1 and anti-human HLA-DR antibody were added in polarization panel. While in Treg panel, anti-CD25 and anti-PD1 was added. After that cells in both the panel was incubated at room temperature for 30 minutes, washed and finally fixation carried out. For the Treg panel, intracellular staining was performed as described above for PBMC using anti-FOXP3 and anti-CTLA4 antibodies. All events were acquired using the BD Fortessa X450 and data analyzed using Flow Jo version 10.6. Flourescent minus one (FMO) controls were included as cut offs for placement of gates when handling nondiscriminatory variables.

### Statistical analysis

Graph Pad Prism 8.0 software was used to carryout statistical comparisons and plot the figures. Descriptive statistics were presented to summarize continuous variables. Wilcoxon rank-sum tests were used to determine the statistically significant difference in continuous outcomes between morphine and saline groups. For flow cytometry data, Mann Whitney T tests were used for unpaired comparisons between the morphine and saline groups. All p values less than 0.05 are considered statistically significant.

## Acknowledgements

We thank staffs and veterinarians of University of Nebraska Medical Center, Comparative Medicine department staffs for housing and animal procedures. We also thank other laboratory members of SNB, SB and HSF for helping in morphine and ART injections to rhesus macaques.

## Authors contributions

AA: Design and performed the experiments, analyze the data, wrote the manuscript; MT: Assist in DNA/RNA purification and ddPCR reaction set-up; OAO: Performed flow experiments, analyze and interpret data. KP: Performed the plasma and CSF viral load measurement experiments; BM: Performed isolation and purification of brain CD11b+ macrophages. BL: Performed brain macrophage purifications and animal experiments. SC: Performed animal experiments. QF: Performed statistical analysis. HSF, SB, SNB: Overall study design, macaque experiments, funding, and edited the manuscript. All authors reviewed the final draft of the manuscript.

## Abbreviations

BAL: Bronchoalveolar lavage
BBB: Blood Brain Barrier
cART: combined Antiretroviral Therapy
CCL2: C-C Motif Chemokine Ligand 2
CCL5: C-C Motif Chemokine Ligand 5
CCR5: C-C chemokine receptor type 5
CD4+: T cells Cluster of differentiation 4+ T cells
CNS: Central Nervous System
CSF: cerebrospinal fluid
ddPCR: Droplet Digital Polymerase Chain Reaction
DMEM: Dulbecco’s Modified Eagle Medium
DNA: Deoxyribonucleic acid
DRV: Darunavir
DTG: Dolutegravir
EDTA: Ethylenediaminetetraacetic acid
FBS: Fetal bovine serum
FTC: Emtricitabin
GALT: Gut Associated Lymphoid Tissues HIV-1 Human Immunodeficiency Virus - 1
HCV: Hepatitis C virus
IACUC: Institutional Animal Care and Use Committee
IBC: Institutional biosafety Committee
ICAM: Intercellular adhesion molecule
IL-6: Interleukin 6
IPDA: Intact proviral DNA assay
IUPM: Infectious units per million
LN: Lymph Node
MØ-QVOA: Macrophage Quantitative Viral Outgrowth Assay
NHPs: Non-human primates
NK: cells Natural killer cells
OUD: Opioid use disorder
PBMCs: Peripheral blood mononuclear cells
PBS: Phosphate-buffered saline
PLWH: People living with HIV
PMA: Phorbol Myristate Acetate
QVOA: Quantitative Viral Outgrowth Assay
RM: Rhesus Macaque
RNA: Ribonucleic acid
RPM: Revolutions per minute
RTV: Ritonavir
SIV: Simian immunodeficiency virus
SNP: Single Nucleotide Polymorphisms
SUD: Substance use disorders
TFV: Tenofovir alafenamide
TILDA: Tat/rev Induced Limiting Dilution Assay
TLR4: Toll-like receptor 4
TNF-α: Tumor necrosis factor alpha
VCAM: Vascular cell adhesion protein
β-ME: 2-Mercaptoethanol

---

**S1 Table:**
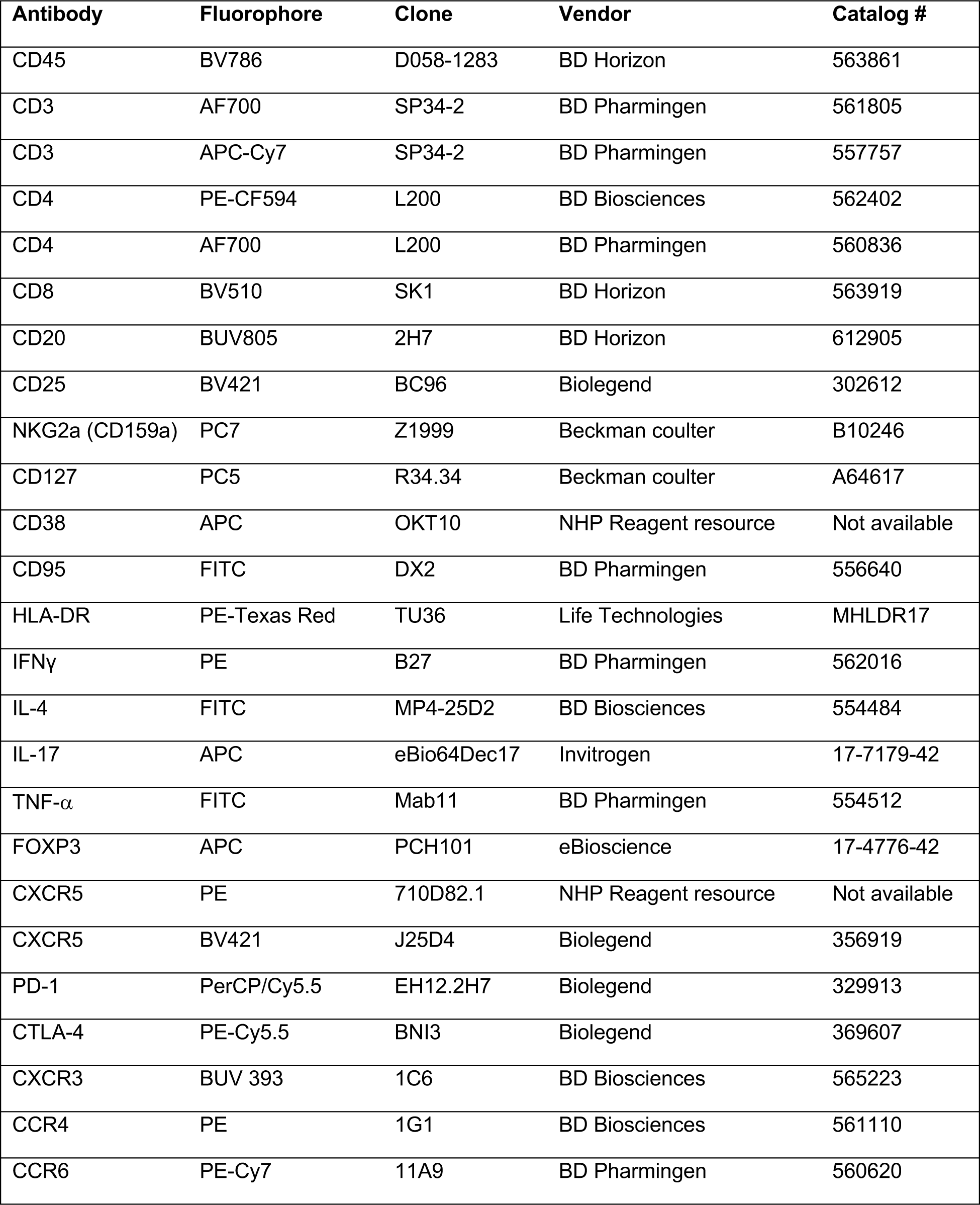
The details of all fluorochrome-conjugated monoclonal antibodies used in the flow cytometry experiments.

| Antibody | Fluorophore | Clone | Vendor | Catalog # |
| --- | --- | --- | --- | --- |
| CD45 | BV786 | D058-1283 | BD Horizon | 563861 |
| CD3 | AF700 | SP34-2 | BD Pharmingen | 561805 |
| CD3 | APC-Cy7 | SP34-2 | BD Pharmingen | 557757 |
| CD4 | PE-CF594 | L200 | BD Biosciences | 562402 |
| CD4 | AF700 | L200 | BD Pharmingen | 560836 |
| CD8 | BV510 | SK1 | BD Horizon | 563919 |
| CD20 | BUV805 | 2H7 | BD Horizon | 612905 |
| CD25 | BV421 | BC96 | Biolegend | 302612 |
| NKG2a (CD159a) | PC7 | Z1999 | Beckman coulter | B10246 |
| CD127 | PC5 | R34.34 | Beckman coulter | A64617 |
| CD38 | APC | OKT10 | NHP Reagent resource | Not available |
| CD95 | FITC | DX2 | BD Pharmingen | 556640 |
| HLA-DR | PE-Texas Red | TU36 | Life Technologies | MHLDR17 |
| IFNγ | PE | B27 | BD Pharmingen | 562016 |
| IL-4 | FITC | MP4-25D2 | BD Biosciences | 554484 |
| IL-17 | APC | eBio64Dec17 | Invitrogen | 17-7179-42 |
| TNF-α | FITC | Mab11 | BD Pharmingen | 554512 |
| FOXP3 | APC | PCH101 | eBioscience | 17-4776-42 |
| CXCR5 | PE | 710D82.1 | NHP Reagent resource | Not available |
| CXCR5 | BV421 | J25D4 | Biolegend | 356919 |
| PD-1 | PerCP/Cy5.5 | EH12.2H7 | Biolegend | 329913 |
| CTLA-4 | PE-Cy5.5 | BNI3 | Biolegend | 369607 |
| CXCR3 | BUV 393 | 1C6 | BD Biosciences | 565223 |
| CCR4 | PE | 1G1 | BD Biosciences | 561110 |
| CCR6 | PE-Cy7 | 11A9 | BD Pharmingen | 560620 |

**S2 Table:** The number of macrophages used for the MØ-QVOA for each animal is described along with the corresponding IUPM values.

| Serial # | Animal_ID | Brain CD11b+ macrophages |  |
| --- | --- | --- | --- |
|  |  | # of cells used in MØ-QVOA | IUPM |
| 1 | CK03 | 1.67E+07 | 0.185 |
| 2 | CK07 | 1.00E+07 | 0.101 |
| 3 | CK40 | 1.67E+07 | 0.144 |
| 4 | CI04 | 1.67E+07 | 0.219 |
| 5 | CI47 | 3.33E+07 | 0.444 |
| 6 | CK22 | 2.33E+06 | 0.602 |
| 7 | CK31 | 1.00E+07 | 1.194 |
| 8 | CK48 | 3.33E+06 | 3.33 |
| 9 | CH89 | 1.67E+07 | 1.569 |
| 10 | CI83 | 1.67E+07 | 0.473 |
| 11 | CI90 | 1.67E+07 | 0.227 |

**S1 Fig:**
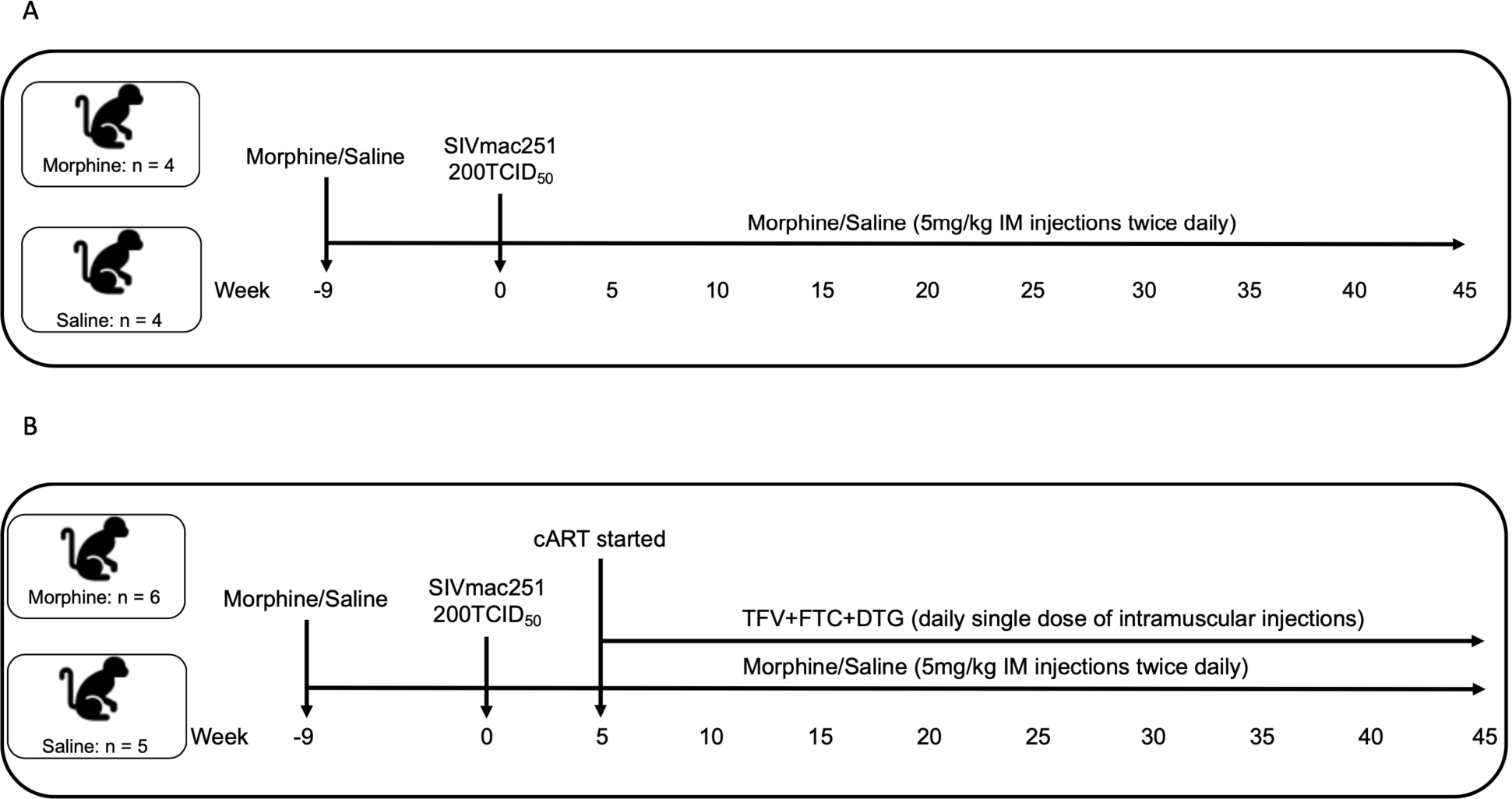
Experimental schema utilized for the study. The study included a total of 19 rhesus macaques. The study had two experimental arms, one of which contain eight monkeys that did not received antiretroviral therapy (A) and the other group had eleven macaques those were treated with combinational antiretroviral therapy (B). (A) In this group out of eight animals four were ramp up with 5 mg/kg intramuscular injection of morphine twice daily for two weeks and then continue morphine dose for seven weeks and the other four animals received similar dose of normal saline (control group). After nine weeks all the macaques were intravenously infected with 200TCID50 of SIVmac251 while the administration of morphine/saline continued until end of the study. (B) In this group out of eleven animals six were ramp up with 5 mg/kg intramuscular injection of morphine twice daily for two weeks and then continue morphine dose for seven weeks and the other five animals received similar dose of normal saline (control group). After nine weeks all the macaques were intravenously infected with 200TCID50 of SIVmac251. Five weeks post infection, anti-retroviral therapy (ART) was initiated and continued until end of the study. ART regimen consisted of two reverse transcriptase inhibitors (FTC: 40 mg/ml and TFV: 20 mg/ml) and one integrase inhibitor (DTG: 2.5 mg/ml). The anti-retroviral drugs were administered subcutaneously once daily at 1 ml/kg body weight, while the administration of morphine/saline continued until end of the study.

**S2 Fig:**
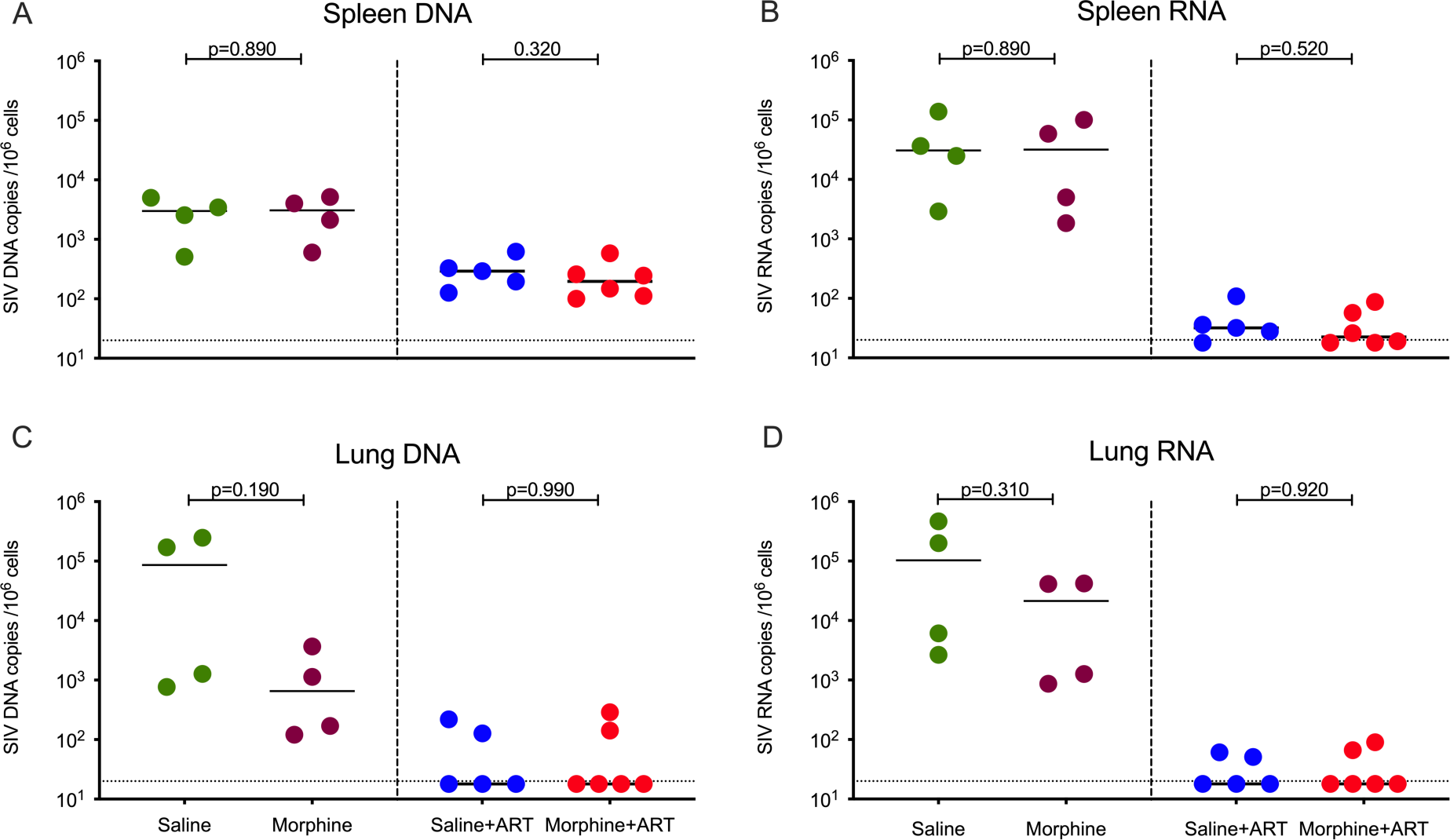
Cell associated SIV DNA and RNA level in spleen and lung in morphine administered versus control group of SIVmac251 infected *rhesus maca*ques. **(A)** SIV DNA copies per million cells from spleen; **(B)** SIV RNA copies per million cells from spleen; (**C)** SIV DNA copies per million alveolar macrophages from lung; **(D)** SIV RNA copies per million alveolar macrophages from lung; The green dots represent samples from control animals administered saline that did not received ART, maroon dots represent samples from morphine administered animals that did not received ART, blue dots represent samples from control animals administered saline and treated with ART and the red dots represent samples from morphine administered animals treated with ART. The horizontal dashed lines represent limit of detection of the assay (20 copies/ml). All values bellow limit of detection are bringing to the limit of detection for better representation in the figure.

**S3 Fig:**
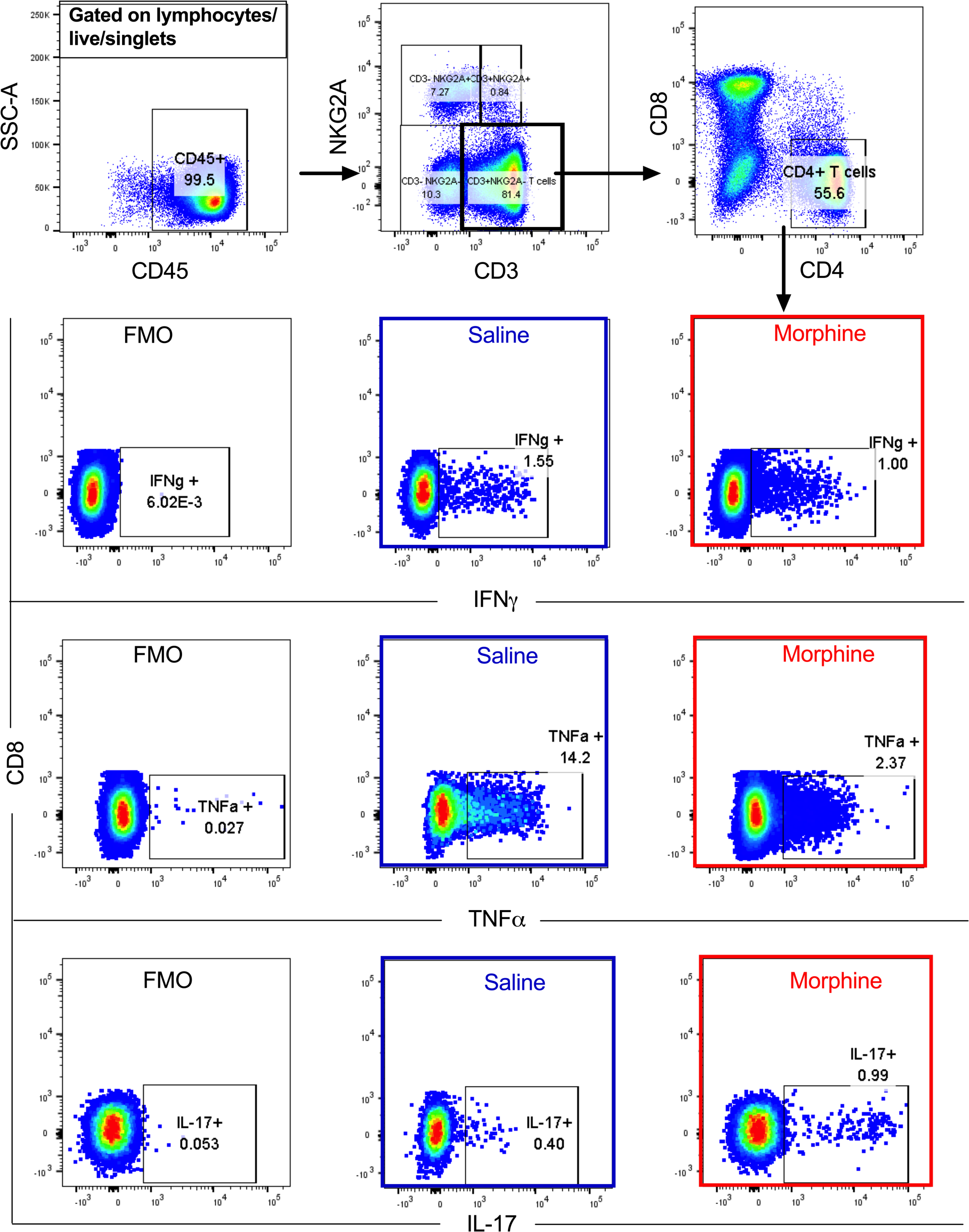
Gating strategy for CD4+ Th1/Th17 polarity in PBMCs. Briefly, CD45+ cells were gated and later NKT cells excluded from the total CD3+ T cell pool. CD4 and CD8 T cells were gated out of the total T cells. The CD4 T cells were then further investigated to determine the extent of IFNγ, TNFα and IL-17 cytokine secretion following PMA and ionomycin stimulation.

**S4 Fig:**
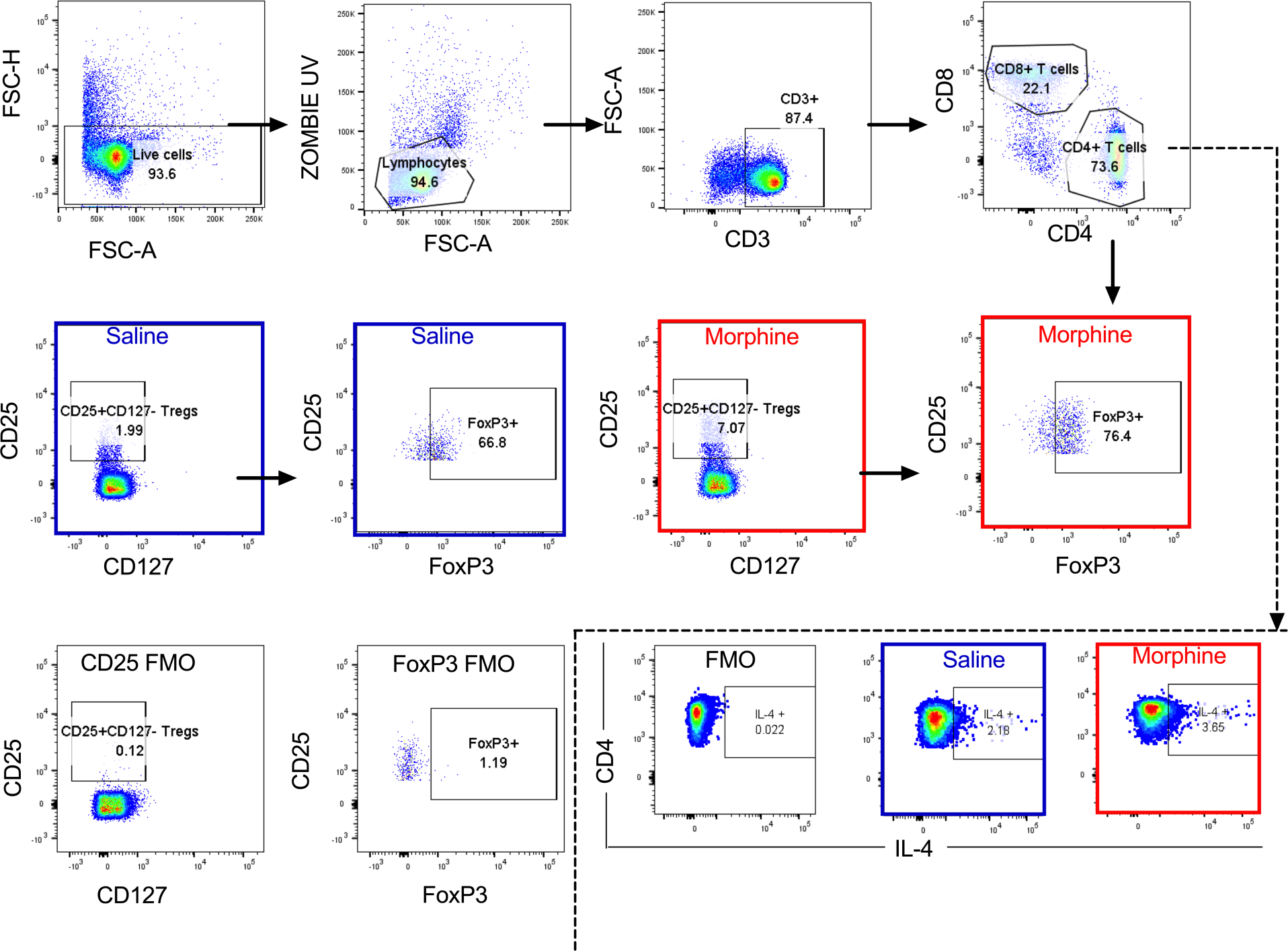
Gating strategy for Th2/T regs in PBMCs. Dead cells excluded based on zombie UV dye positive expression. Based on FSC-A vs SSC-A gating, lymphocytes were later gated and CD4+ T cells further obtained from total CD3+ T cells. From the CD4+ T cells, the extent of IL-4 cytokine secretion was then evaluated following stimulation with PMA/ ionomycin. Finally, the frequencies of CD4+ T regs were evaluated based on the expression of CD25+ and CD127-. These cells were later evaluated for their expression of the transcription factor of FoxP3+ to give the final frequency of T regs denoted as %CD4+ CD25+ CD127-Foxp3+ cells.

**S5 Fig:**
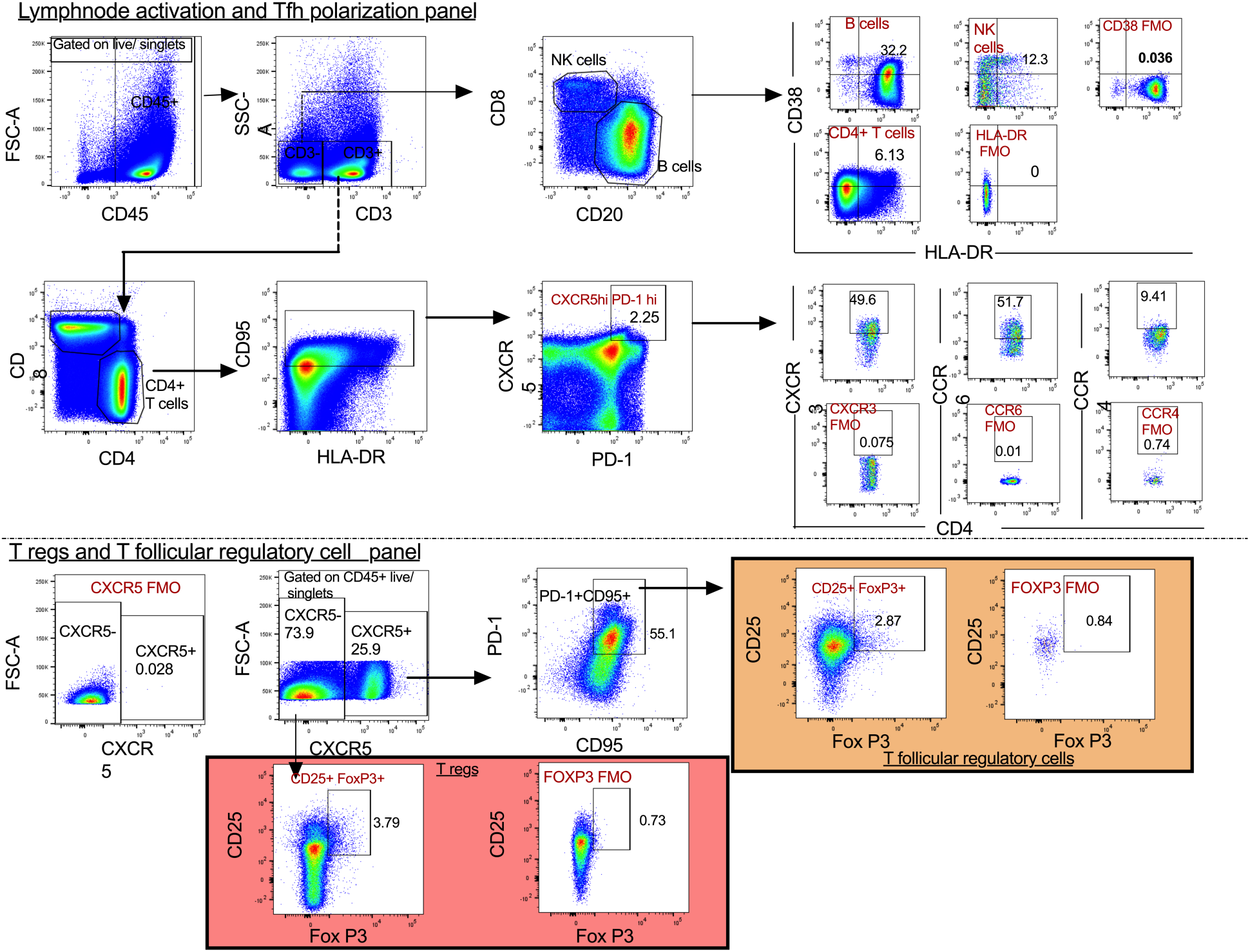
Gating strategy for Lymph node activation and Tfh Polarization panel. CD45+ leukocytes were gated and T and non-T lymphocytes discriminated based on CD3+ expression. CD3-cells were segregated based on CD20+ (B cells) and CD8α+ (NK cells). The activation profiles of these cells were further evaluated based on CD38+ and HLA-DR+ co-expression. CD4+ T cells were separately gated out of CD3+ T cells and levels of activation determined. The CD95+ memory cell marker was then included to obtain memory CD4+ T cells and later delineate the Tfh population based on CXCR5 and PD-1. Additional chemokines were used to study Tfh polarity based on Tfh1 (CXCR3), Tfh2 (CCR4), Tfh17 (CCR6) and T follicular regulatory cells (CD25 and FoxP3).

**S6 Fig:**
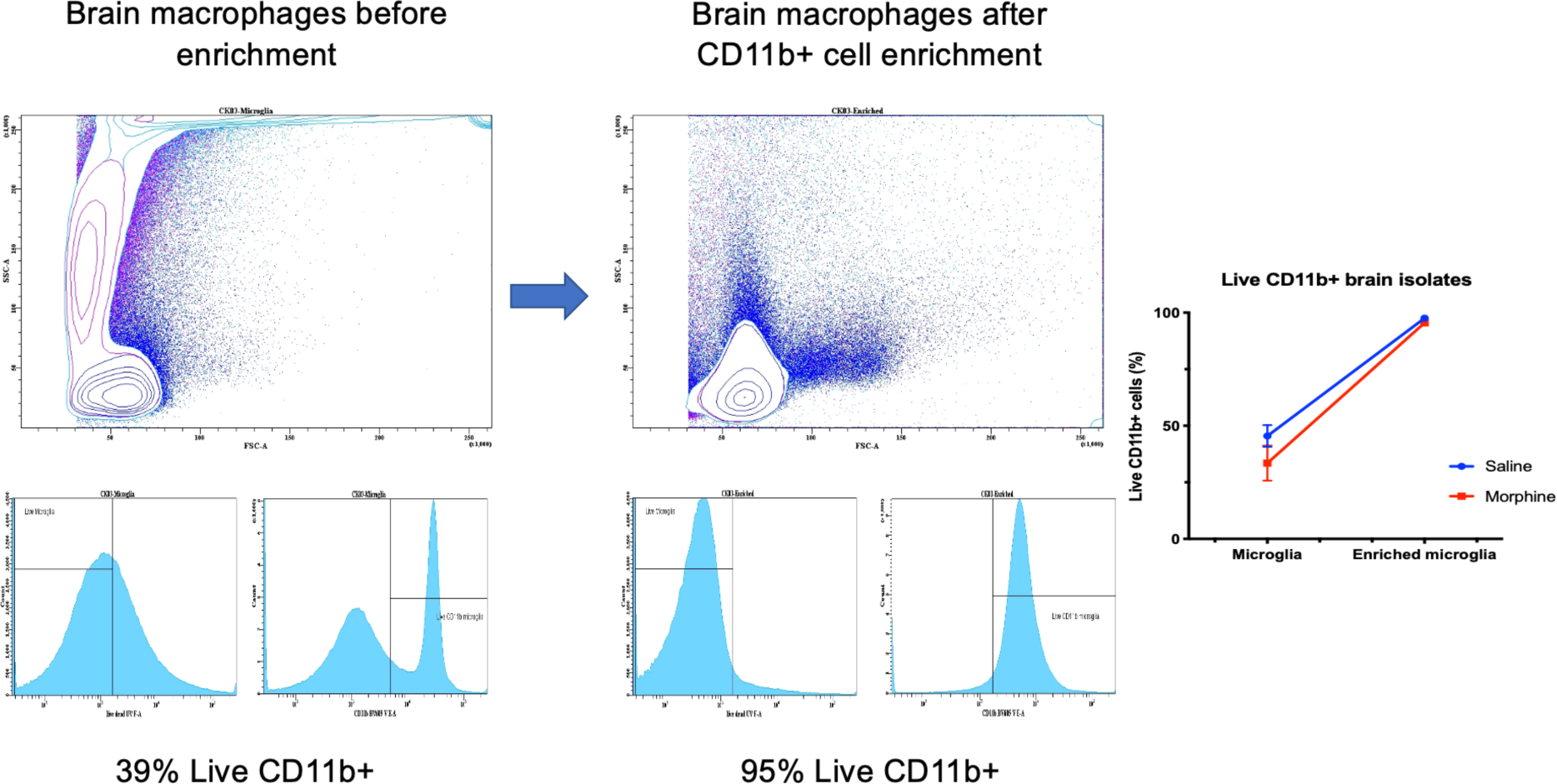
Gating strategy to detect the purity of CD11b+ macrophages isolated from brain of rhesus macaques as described. Briefly, comparisons in the subsequent purity of CD11B microglia showed that higher yields of brain macrophages were obtained following enrichment of brain cells with CD11b+ beads.

